# Developmental rewiring of the *NGAL*/*CUC*/*KLU* network associated with pleiotropic roles of *NGAL* genes

**DOI:** 10.1101/2024.12.12.627899

**Authors:** Antoine Nicolas, Panagiotis Papadopoulos, Mattéo Caroulle, Bernard Adroher, Magali Goussot, Anne-Sophie Sarthou, Nicolas Arnaud, Aude Maugarny, Patrick Laufs

## Abstract

Gene Regulatory Networks (GRNs) play prominent roles in regulating developmental processes, and their modulation across species is a major source for evolutionary innovation. However, it remains poorly understood how GRNs are rewired between different organs within a single species. This question is particularly relevant for pleiotropic genes, which may exhibit organ-specific GRN modulations potentially reflecting their diverse functions. To address this, we investigated the *NGATHA-like* (*NGAL*) genes, as a model for pleiotropic genes that regulate growth or patterning in multiple Arabidopsis organs via two distinct pathways involving the *CUP-SHAPED COTYLEDON* (*CUC*) and *KLUH* (*KLU*) genes. By combining genetic analysis with gene expression characterization, we uncovered significant organ-specific rewiring of the *NGAL*/*CUC*/*KLU* regulatory module. Our findings highlight that changes in gene expression patterns, potentially arising from developmental constraints, play a pivotal role in the organ-specific modulation of GRNs. Furthermore, GRNs at the molecular and functional levels do not always align perfectly, potentially due to the influence of additional regulatory mechanisms. Altogether, our findings reveal significant modulation of the GRNs associated with pleiotropic genes. We propose that this flexibility in GRNs facilitates gene pleiotropy.

## Introduction

A concept that has emerged during the two last decades is that biological processes are governed by gene regulatory networks (GRNs) (Benfey and Weigel 2001; Davidson et al. 2003). While examining a GRN in a specific species or organ provides valuable insights into how it controls development and physiology, comparing GRNs across different species offers diverse insights into evolutionary processes. Thus, investigating the conservation of GRNs across species is a way to explore homology between structures in distantly related species, while GRN modulations can shed light on how innovations may have occurred during evolution (Doebley and Lukens 1998; Hinman et al. 2003; Scotland 2010; Lynch and Roth 2011; Peter and Davidson 2011; Das Gupta and Tsiantis 2018; Cary et al. 2020; Srivastava 2021; Swift et al. 2024). In parallel, the more recent integration of single-cell transcriptomic data with other genome-wide omics has started to provide an unprecedent view of the GRN diversity at the cellular and tissular level (Qian and Huang 2020; Longo et al. 2021; Badia-i-Mompel et al. 2023). However, understanding the functional significance of such GRN variations remains challenging. Here, we genetically addressed the question of inter-organ modulation using as a model a GRN that regulates multiple aspects of aerial organ development in Arabidopsis.

A major mechanism driving the evolution of GRN structure and/or function is gene co-option (Halfon 2017; Jiggins et al. 2017; Das Gupta and Tsiantis 2018). Gene co-option occurs when changes in the *cis*-regulatory sequences of a gene cause it to become active in a novel developmental context where it may acquire a novel function. Because genes do not function in isolation, a resulting question is to which extent the GRN associated with the co-opted gene is remodeled in the new biological environment (Tyler et al. 2009; McQueen and Rebeiz 2020). Indeed, two opposite evolutionary patterns of GRN rearrangements have been observed. First, GRNs can keep their overall topology despite being redeployed in distantly related organisms. For instance, a similar GRN is promoting suberin deposition in the Arabidopsis endodermis and the tomato exodermis, two cell layers that share similar biological roles despite different positions in the root (Cantó-Pastor et al. 2024). Redeployment of a conserved GRN can be also associated with different functions, as for instance a conserved GRN that controls eye development in the fly and cell migration in the sea urchin (Martik and McClay 2015). Second, morphological innovations during evolution can be driven by co-opted GRNs that are partially modified such as those for the formation of trichomes or related unicellular projections in different organs or species of Drosophila (Kittelmann et al. 2018; Rice et al. 2024), or that are more profoundly rewired, as shown for the development of simple and compound leaves (Rast-Somssich et al. 2015)

Another way to address GRN remodeling is to compare GRNs between different organs within a species. This is particularly relevant for GRNs centered around pleiotropic genes. Pleiotropy is defined as a phenomenon where a single gene or locus influences two or more distinct phenotypic traits (Stearns 2010; Paaby and Rockman 2013). Remodeling of some GRNs between organs has been suggested in plants. For instance, the WUSCHEL/CLAVATA (WUS/CLV) network shows a regulation by AGAMOUS which is specific to the floral meristem and absent from the shoot apical meristem (Lenhard et al. 2001). One can however argue that in this particular example the WUS/CLV function is not truly pleiotropic between floral and apical meristems, as in both organs it regulates stem cell homeostasis (Schoof et al. 2000; Lenhard et al. 2001). The *AINTEGUMENTA-LIKE/PLETHORA* genes provide another example of organ-dependent rewiring of the GRN as these genes activate cell wall remodeling genes in the shoot while repressing them in the root (Scheres and Krizek 2018), but yet again the pleiotropy of these genes could be discussed as they promote growth in both organs. Here, we investigate the remodeling of a small GRN centered on the pleiotropic *NGATHA-like* (NGAL) genes in different Arabidopsis organs.

The *NGATHA-like*, *NGATHA-like1* (*NGAL1*)/*ABNORMAL SHOOT 2* (*ABS2*), *NGAL2/SUPPRESSOR OF DA1 7* (*SOD7*) and *NGAL3/DEVELOPMENT-RELATED PCG TARGET IN THE APEX 4* (*DPA4*) genes encode transcription factors of the B3 family (Swaminathan et al. 2008; Romanel et al. 2009). Analysis of *ngal* simple and double mutants have revealed pleiotropic roles for these genes during the development of the aerial organs (Engelhorn et al. 2012; Shao et al. 2012, 2020; Chen et al. 2019; Nicolas et al. 2022; Zheng et al. 2023). For instance, *DPA4* and *SOD7* redundantly limit growth, particularly in seed, cotyledon and floral organs as these organs are larger in the *dpa4 sod7* double mutant (Zhang et al. 2015; Chen et al. 2019; Zheng et al. 2023). They also contribute to the establishment of phyllotaxis and to the development of axillary meristems. The *dpa4 sod7* double mutant shows an altered phyllotaxis with the appearance of flower clusters on the main shoot and delayed axillary meristem initiation and outgrowth (Chen et al. 2019; Nicolas et al. 2022). Finally, *DPA4*, *SOD7* and *ABS2* limit the extent of leaf margin serration (Shao et al. 2020).

The pleiotropic role of *NGAL* genes during development has been associated with the function of two genetic pathways. First, there is strong evidence for genetic and molecular interactions linking *NGAL* and *CUP-SHAPED COTYLEDON* (*CUC*) genes, which code for NAC-family transcription factors (Aida et al. 1997; Takada et al. 2001; Vroemen et al. 2003). Thus, interactions between *NGAL* and *CUC* genes shape leaf margins. The *dpa4-1 sod7-1 abs2-2* triple mutant shows increased leaf margin serration compared to a wild-type plant, thus resembling plants with increased *CUC* activity (Nikovics et al. 2006; Shao et al. 2020). On the contrary, lines overexpressing *ABS2*/*NGAL1* or *DPA4*/*NGAL3* have reduced leaf serration, similar to *cuc2* or *cuc3* mutants (Nikovics et al. 2006; Engelhorn et al. 2012; Shao et al. 2012, 2020). A similar interaction between *NGAL* and *CUC* genes occurs during axillary meristem development. *CUC* genes are required for axillary meristem initiation, likely by preventing cell differentiation and maintaining cells in a meristematic fate (Raman et al. 2008). However, *CUC* genes need to be repressed by *DPA4* and *SOD7* for the initiating meristem to proceed to the establishment phase and allow stem cell formation and proper expression of *WUS* and *CLV3* (Nicolas et al. 2022). Several lines of evidence suggest the control of leaf and axillary meristem development by the *NGAL* genes mostly occurs via their repression of the *CUC* genes. First, NGAL proteins can bind to *CUC* promoters, and in *ngal* mutants the expression of the *CUC* genes is increased and/or ectopic. Second, transcriptomic data show that NGAL-mediated gene expression regulation is largely dependent on *CUCs* (Shao et al. 2020). Third, leaf and axillary meristem defects of *ngal* mutants are mostly suppressed when *CUC* genes are additionally mutated (Shao et al. 2020; Nicolas et al. 2022).

The second pathway acting downstream the *NGAL* genes involves the *KLUH* (*KLU*) gene which codes for a cytochrome P450 of the CYP78A subfamily (Anastasiou et al. 2007). KLU promotes growth in a non-cell-autonomous manner, possibly through a yet-to-identify mobile signal that maintains cell proliferation (Anastasiou et al. 2007; Adamski et al. 2009; Eriksson et al. 2010; Kazama et al. 2010). Genetic analyses show that *DPA4* and *SOD7* act in a common pathway with *KLU* to regulate seed size (Zhang et al. 2015). Thus, a lack of *KLU* repression by the DPA4 and SOD7 proteins lead to larger seeds as observed in the *dpa4-2 sod7-3* mutant. In addition, SOD7 directly binds to the *KLU* promoter to repress its expression (Zhang et al. 2015).

Taken together, these data are demonstrating a strong genetic link between both *CUC* and *NGAL* genes and *KLU* and *NGAL* genes. Recent data also suggest a genetic link between the *CUC* and *KLU* genes. During embryo development and leaf margin morphogenesis, expression patterns of the *CUC* and *KLU* genes overlap (Maugarny-Calès et al. 2019; Aida et al. 2020). Furthermore, *KLU* expression is absent at these two stages in *cuc* mutants, while its expression is induced following *CUC* activation. This suggests that *KLU* is acting downstream of the *CUC* genes. Interestingly, while expressing *KLU* under the control of *CUC* regulatory sequences is not sufficient to restore embryo defects of *cuc* mutants, it is to promote leaf serration (Maugarny-Calès et al. 2019; Aida et al. 2020). This indicates that *KLU* has a more prominent role in the network acting downstream of *CUC* genes during leaf development than it has during embryo development.

These elements indicate that the *NGAL* genes are important factors controlling plant development in a pleiotropic way and that this involves two possible pathways, involving either *CUCs* or *KLU* genes. However, we lack a systematic analysis of the contribution of these two pathways in mediating the function of the NGAL transcription factors in different organs, which could reveal possible organ-specific modulation of the *NGAL* GRN. That such modulation indeed exists is suggested by the observation that *NGAL* genes, despite being expressed during both embryonic and post-embryonnic meristem formation, are only required for the latter one (Nicolas and Laufs 2022; Nicolas et al. 2022). Therefore, we explored here whether the *NGAL*/*KLU*/*CUC* module was modified between different organs. We first show that within the three *NGAL* genes, *DPA4* and *SOD7* have the most prominent role and redundantly control several aspects of aerial development, with, however, small variations in the contributions of the different genes. Next using a genetic approach, we reveal inter-organ modulation of the *NGAL*/*KLU*/*CUC* module. Hence while *NGALs* repress floral organ growth mostly via the *KLU* pathway, their repressive effect on cauline leaf growth occurs through both *KLU* and *CUC* pathways. Distinct from these, *NGAL* controls sepal numbers and phyllotaxis via *CUC* genes and mostly independently of *KLU*. Finally, *NGAL* and *CUC* independently repress seed growth. We further show that such network modulation can be due to different expression patterns of the acting genes. We also show that genetic regulations between components of the network are not necessarily translated into a functional effect, showing that molecular and functional networks may differ in an organ-specific way. Altogether, we provide evidence for the rewiring of the GRNs between different plant organs, and we suggest that such a flexibility of the GRN facilitates gene pleiotropy.

## Results

### *DPA4* and *SOD7* are the two major *NGAL* genes redundantly controlling plant development

Previous reports indicate that *dpa4 sod7* double mutants (*ds*) are affected in multiple aspects of aerial organ development as they show larger cotyledons, flowers and seeds, exhibit increased leaf serration and have phyllotaxy and branching defects (Engelhorn et al. 2012; Shao et al. 2012, 2020; Chen et al. 2019; Nicolas et al. 2022; Zheng et al. 2023). However, apart from shoot branching (Nicolas et al. 2022), the contribution of individual *NGAL* genes to these phenotypes and their genetic redundancy was not systematically addressed. Therefore, we quantified floral organ numbers, the size of cauline leaves, petals, pistils and seeds and the phyllotaxis in the three single and three double *ngal* mutants (Figure 1, Figure S1). All these parameters were significantly different in *ds* compared to the wild type (WT) (Figure 1, Figure S1). Cauline leaves, petals, pistils and seeds were larger in *ds* compared to WT (Figure 1A, G, I). More flowers were located in clusters along the stem (Figure 1C). The *ds* flowers had a higher average number of sepals, petals, and carpels, but a lower average number of stamens (Figure 1E, Figure S1A-H). In contrast, single mutants and the two other double mutant combinations (*abs2-1 dpa4-2* and *abs2-1 sod7-2*) showed only limited phenotypic abnormalities. Only the mean number of sepals was significantly increased in the *abs2-1 sod7-2* double (Figure 1E). However, the number of sepals, petals and/or stamens were different from those in the WT in about 10% to 30% of *ds* flowers (Figure S1E-H), suggesting *NGAL* genes contribute to floral phenotype robustness. In addition, cauline leaves were slightly larger in *dpa4-2* and *abs2-1 sod7-2,* as were pistils in *abs2-1 dpa4-2* and *abs2-1 sod7-2* (Figure 1A). In contrast, petals were smaller in *abs2-1* (Figure S1I). Together, these observations indicated that each *NGAL* gene has only a very limited unique role. In contrast, *DPA4* and *SOD7* are mostly redundant in their roles to pleiotropically control plant development. This suggests that these two genes may have undergone limited subfunctionalization and neofunctionalization since their recent formation by duplication around 24 to 40 million years ago (Romanel et al. 2009). Next, to genetically determine the contributions of the *KLU* and *CUC* pathways to the pleiotropic roles of *DPA4* and *SOD7*, we analyzed the phenotype of *ds* mutants in which *KLU* or *CUC* genes were inactivated.

**Figure 1.**
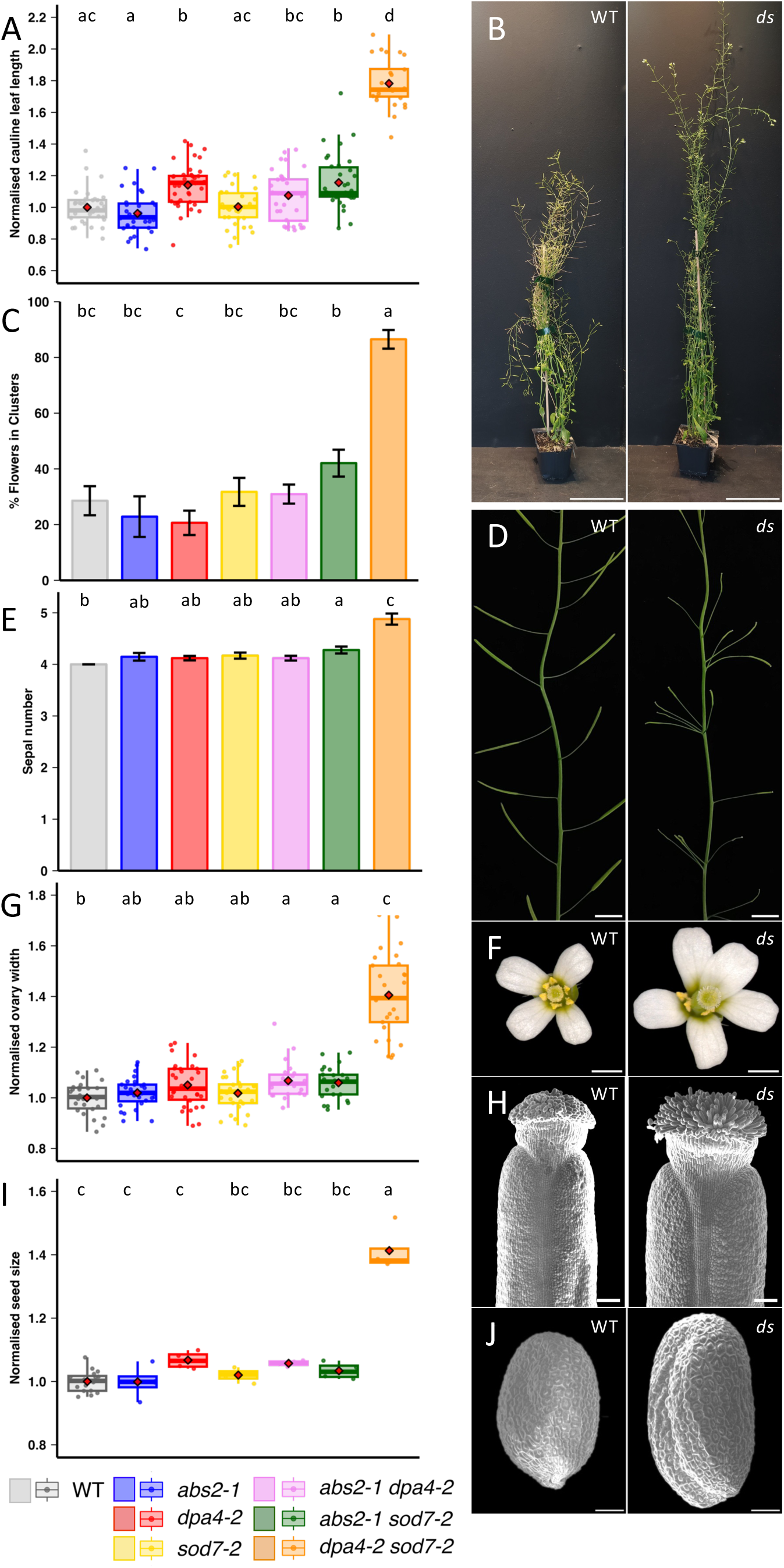
*DPA4* and *SOD7* have redundant, pleiotropic roles during plant development. **A.** Cauline leaf length in WT, single and double *ngal* mutants. **B.** Mature wild-type (WT) and *dpa4-2 sod7-2* double mutant (*ds*) plants. **C.** Percentage of flowers grouped in clusters in WT, single and double *ngal* mutants. **D.** Part of the main inflorescence stem of wild-type (WT) and *dpa4-2 sod7-2* double mutant (*ds*) plants. **E.** Sepal number in WT, single and double *ngal* mutants. **F.** Top view of mature wild-type (WT) and *dpa4-2* and *sod7-2* double mutant (*ds*) flower. **G.** Ovary width in WT, single and double *ngal* mutants. **H.** Scanning electron microscopy detail of wild-type (WT) and *dpa4-2 sod7-2* double mutant (*ds*) pistil top part. **I.** Seed size in WT, single and double *ngal* mutants. **J.** Scanning electron microscopy view of wild-type (WT) and *dpa4-2 sod7-2* double mutant (*ds*) seeds. In **A**, **G** and **I** data were normalized to the mean value of the WT. Rectangle represent the first and third quartiles, with a horizontal central line marking the median. Mean values are represented as red diamonds. In **A**, **E** and **G,** a Kruskal-Wallis test followed by a Dunn’s post-hoc test were performed to show significant differences between different mutants and in **C** and **I,** ANOVA followed by a Tukey post-hoc test were performed (p-value<0.05). Scale bars = 10 cm in A, 1 cm in D, 1 mm in F, 100 µm in H and J.

### *DPA4* and *SOD7* redundantly repress floral organ growth via *KLU,* independently of *CUC* genes

The increase in petal and pistil size observed in the *ds* mutants was not affected by additional mutation of any of the *CUC* genes (Figure 2A, B, Figure S2A, B). Furthermore, petals and pistils sizes were not modified in *cuc* single mutants, with the exception of a reduced size of *cuc2-1* petals revealing a role of *CUC2* in petal growth. In contrast, *klu-4* mutants developed smaller petals (Figure 2A, C, Anastasiou et al. 2007; Zhang et al. 2015) and pistils (Figure S2A, C). In *dpa4-2 sod7-2 klu-4* triple mutants, petal size was fully restored (Figure 2A, C, Zhang et al. 2015), while pistil size was only partially restored (Figure S2A, C). Thus, *DPA4* and *SOD7* repress floral organ growth via *KLU* independently of *CUC* genes (Figure 6A). However, the observation that the *dpa4-2 sod7-2 klu-4* triple mutants has petals and pistils larger than *klu-4*, suggest that *DPA4* and *SOD7* also repress growth via another pathway independent of *KLU*. Additionally, the partial restoration of pistil size in *dpa4-2 sod7-2 klu-4* triple mutants also supports this hypothesis (Figure 6A).

**Figure 2.**
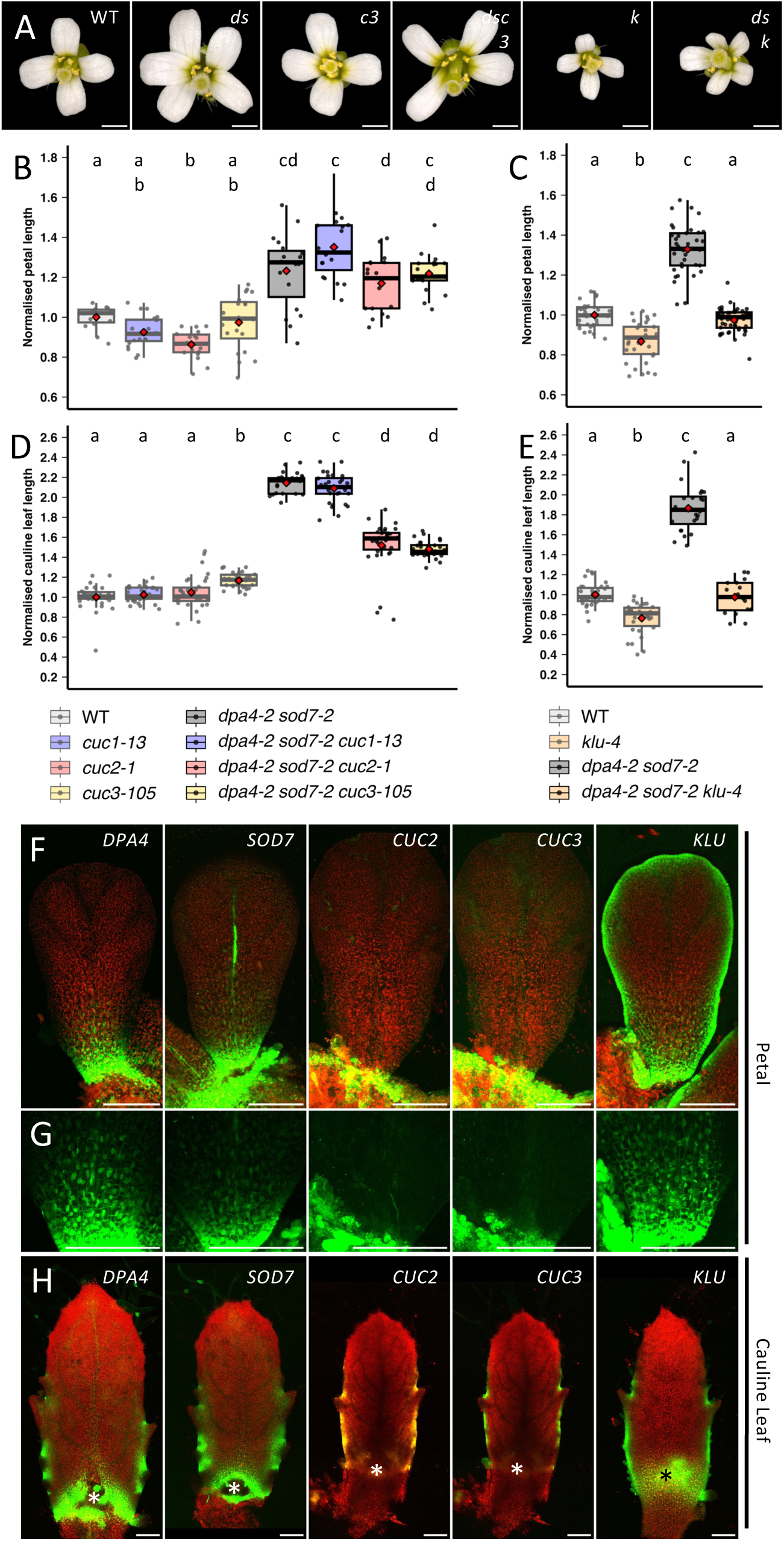
*DPA4*/*SOD7* redundantly repress petal growth via *KLU* and cauline leaf growth via *KLU* and *CUC* genes. **A.** Top view of mature wild type (WT), *dpa4-2 sod7-2* (*ds*), *cuc3-105* (*c3*), *dpa4-2 sod7-2 cuc3-105* (*dsc3*), *klu-4* (*k*) and *dpa4-2 sod7-2 klu-4* (*dsk*) flower. **B.** Petal length in WT, single *cuc*, *dpa4-2 sod7-2* double and *dpa4-2 sod7-2 cuc* triple mutants. **C.** Petal length in WT, single *klu-4*, *dpa4-2 sod7-2* double and *dpa4-2 sod7-2 klu-4* triple mutants. **D.** Cauline leaf length in WT, single *cuc*, *dpa4-2 sod7-2* double and *dpa4-2 sod7-2 cuc* triple mutants. **E.** Cauline leaf length in WT, single *klu-4*, *dpa4-2 sod7-2* double and *dpa4-2 sod7-2 klu-4* triple mutants. **F.** Expression pattern in young developing petals of *pDPA4:GFP* (*DPA4*), *pSOD7:GFP* (*SOD7*), *pCUC2:RFP* (*CUC2*), *pCUC3:CFP* (*CUC3*) and *pKLU:GFP* (*KLU*) reporters. **G.** Detail of **F**, showing the expression of the reporters in the lower part of the petals. **H.** Expression pattern in young developing cauline leaves of *pDPA4:GFP* (*DPA4*), *pSOD7:GFP* (*SOD7*), *pCUC2:RFP* (*CUC2*), *pCUC3:CFP* (*CUC3*) and *pKLU:GFP* (*KLU*) reporters. For **F** to **H**, all reporters are represented in green regardless of the fluorescent reporter used. In **F** and **H**, chlorophyll fluorescence is shown in red. Note that the pictures of *pCUC2:RFP* (*CUC2*), *pCUC3:CFP* (*CUC3*) are from a same sample expressing both reporters simultaneously. In **H**, asterisks mark axillary meristems. In **B** to **E**, data were normalized to the mean value of the WT. Rectangle represent the first and third quartiles, with a horizontal central line marking the median. Mean values are represented as red diamonds. In **B-D,** a Kruskal-Wallis test followed by a Dunn’s post-hoc test were performed to show significant differences between different mutants (p- value<0.05). Scale bars = 100 µm in A, F-H

### *DPA4* and *SOD7* redundantly repress cauline leaf growth via *KLU* and *CUC* genes

Cauline leaves of the *ds* mutants were larger than WT, and this phenotype was partially suppressed by *cuc2-1* or *cuc3-105* mutations and was totally suppressed by the *klu-4* mutation (Figure 2D, E). Mutation of *CUC1* had no effect on cauline leaf size in agreement with the absence of expression of this gene in developing leaves (Hasson et al. 2011). These observations indicated that both *KLU* and *CUC2*/*CUC3* pathways are required for the increased cauline leaf size of the *ds* mutant, though with *CUC2*/*CUC3* having a less important contribution than *KLU* (Figure 6B). Since CUC2 promotes *KLU* expression during leaf development (Maugarny-Calès et al. 2019), *CUC2*/*CUC3* may promote cauline leaf growth via *KLU* activation.

### Rewiring of the *NGAL/CUC/KLU* network between cauline leaves and petals is associated with modified gene expression patterns

Our genetic analyses suggested that the networks controlling floral organ and cauline leaf growth shared a common DPA4/SOD7 – KLU edge but differed by a DPA4/SOD7 – CUC2/3 edge which is specific to cauline leaves (Figure 6A,B). To confirm this, we examined the expression of these genes during petal and cauline leaf development using transcriptional reporters (Figure 2F-H). *DPA4* and *SOD7* transcriptional reporters are strongly expressed in the region where the petals are attached to the flower receptacle and a weaker expression was observed in the proximal base of the petal. In contrast, *CUC2* and *CUC3* reporters where only expressed in the region where the petals are attached to the flower. The *KLU* transcriptional reporter is expressed along the petal margin and in the basal part of the petal (Figure 2F-G). Therefore, *DPA4*, *SOD7* and *KLU*, but not *CUC2* and *CUC3* have overlapping expression patterns in the basal part of the petal, a region in which *KLU* activity has been proposed to regulate organ growth (Kazama et al. 2010). In developing leaves, *DPA4*, *SOD7*, *KLU*, *CUC2* and *CUC3* transcriptional reporters showed overlapping expression pattern along the proximal part of the leaf margin, the expression of the reporters being absent from the developing teeth (Figure 2H), in agreement with the proposed interactions of these genes in cauline leaves (Figure 6B). Therefore, the difference between the GRN regulating cauline leaf and petal growth (Figure 6A,B) may result from the *CUC* genes not being expressed in the latter organ. As strong epidermal auxin response foci have been associated with teeth formation and *CUC* gene expression at the leaf margin (Hay et al. 2006; Kawamura et al. 2010; Bilsborough et al. 2011) we next compared *DR5:VENUS* reporter expression in petal and cauline leaves. This revealed the presence of epidermal foci with strong auxin response present at the margin of cauline leaves which were absent in petals (Figure S2D). Thus, different auxin response pattern may contribute to differential *CUC* expression, which in turn leads to variations in GRNs between petals and cauline leaves. Together, these data indicate that rewiring of the GRN may be the consequence of different expression patterns of the GRN actors.

### *DPA4* and *SOD7* control phyllotaxy via *CUC* gene repression independently of *KLU*

To quantify more precisely the phyllotaxy defects observed in *ds* (Figure 1C, D), we analyzed the distribution of the internode lengths along the main stem. In the *ds* mutant the presence of numerous flower clusters resulting from short internodes separated by long internodes translated in a broader distribution of internode length and a reduction in the median internode length compared to WT (Figure 3A, C). Inactivating individual *CUC* genes in *ds* led to a partial restoration towards a WT phenotype with fewer visible flower clusters (Figure 3C). Quantification confirmed this, as a more compact distribution of the internode length and an increase in the median internode length was observed, with *CUC3* inactivation leading to the best phenotypic restoration (Figure 3A). We could not analyze the phyllotaxy of *ds* with two simultaneous *cuc* mutations because these quadruple mutants showed fusion of the floral pedicel with the main stem. On the other hand, introducing a *klu-4* mutation in *ds* increased its phyllotaxy defects (Figure 3B,C). These observations indicated that the *DPA4*/*SOD7* pathway controls phyllotaxy to a large extent via the *CUC* genes while this pathway is independent of the *KLU* gene (Figure 6D). Phyllotaxy defects of the *ds* mutant were already present early in development, as clusters of flowers were visible at the *ds* inflorescence apex, while in the WT flowers were inserted in a more regular manner (Figure 3D). *CUC2* and *CUC3* transcriptional reporters were ectopically expressed in the young *ds* inflorescence stem, either in large domains or small groups of cells (Figure 3E). Together, these observations indicate that *DPA4*/*SOD7* are required to establish a proper phyllotaxy by repressing ectopic *CUC* expression in developing stem, a similar role as the one proposed for microRNA miR164 that represses *CUC2* (Peaucelle et al. 2007; Sieber et al. 2007).

**Figure 3.**
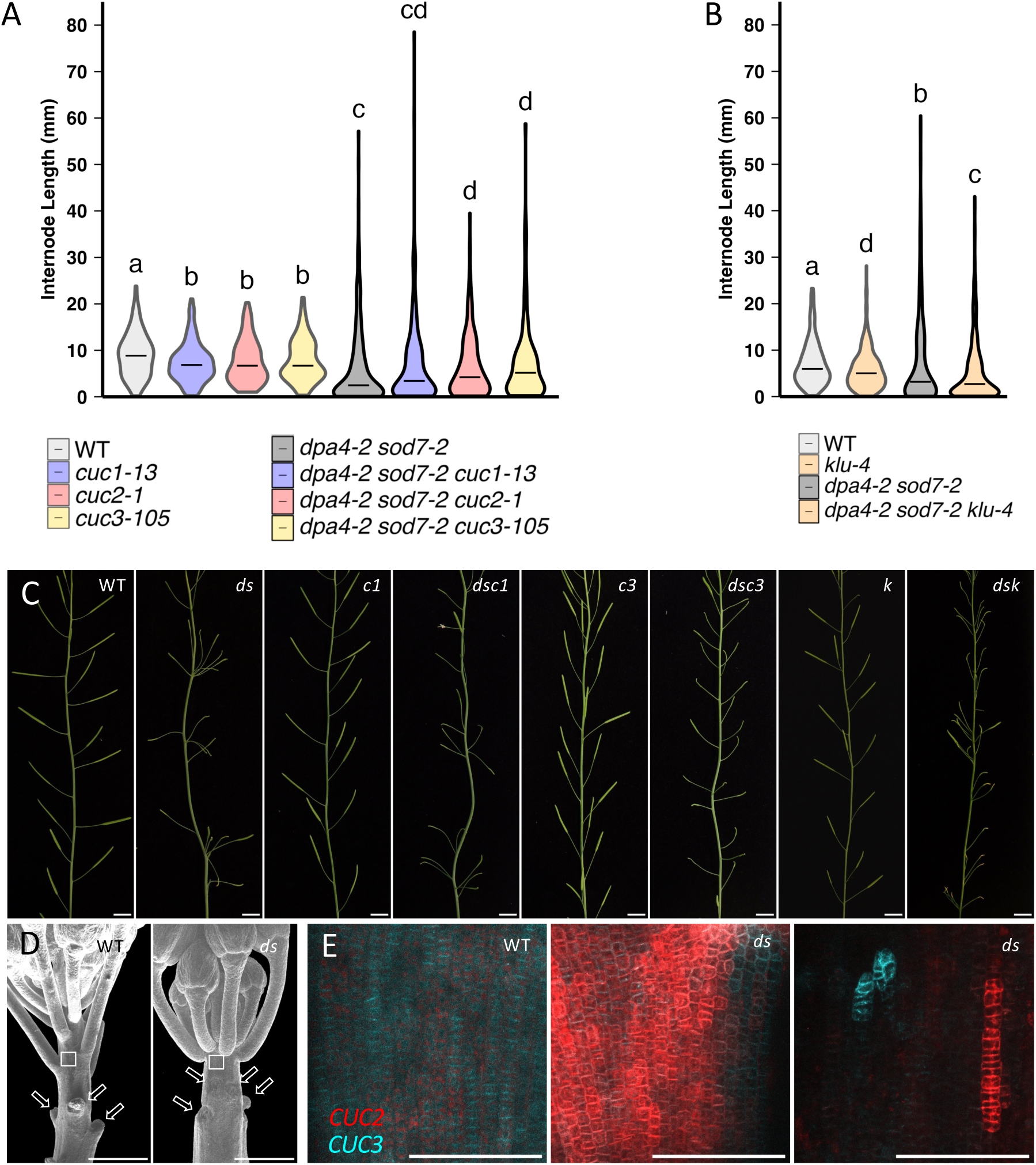
*DPA4*/*SOD7* control phyllotaxis through the *CUC* genes. **A.** Internode length distribution in WT, single *cuc*, *dpa4-2 sod7-2* double and *dpa4-2 sod7-2 cuc* triple mutants. The black horizontal segment represents the median. **B.** Internode length distribution in WT, single *klu-4*, *dpa4-2 sod7-2* double and *dpa4-2 sod7-2 klu-4* triple mutants. The black horizontal segment represents the median. **C.** Part of the main inflorescence stem of wild-type (WT), *dpa4-2 sod7-2* (*ds*), *cuc1-13* (*c1*), *dpa4-2 sod7-2 cuc1-13* (*dsc1*), *cuc3-105* (*c3*), *dpa4-2 sod7-2 cuc3-105* (*dsc3*), *klu-4* (*k*) and *dpa4-2 sod7-2 klu-4* (*dsk*) plants. **D.** Partially dissected inflorescence apex of wild type (WT), *dpa4-2 sod7-2* (*ds*). The flowers were removed from the lower part (arrows). The squares are positioned on the type of zones which were imaged in E. **E.** Expression of *pCUC2:RFP* (in red) and *pCUC3:CFP* (in cyan) in young internodes of wild-type (WT) or *dpa4-2 sod7-2* double mutant (*ds*) plants. In **A,B**, a two sample Kolmogorov-Smirnov test was performed to show significant differences between different mutants (p-value<0.01). Scale bars = 1 mm in D and 100 µm in E.

### *DPA4* and *SOD7* controls floral organ numbers via the *CUC* genes and independently of *KLU*

We next investigated the contribution of the *CUC* and *KLU* genes to the increased sepal and petal numbers observed in *ds* (Figure 1E, F, Figure S1A,B,E,F). The increase in sepal number in *ds* was not modified by any of the single mutation in the *CUC* genes, while mutating two *CUC* genes simultaneously significantly restored the phenotype (Figure 4A, B, Figure S3D,E). Notably the *dpa4-2 sod7-2 cuc1-13 cuc3-105* quadruple mutant had always 4 sepals like the WT (Figure 4B, Figure S3E).

**Figure 4.**
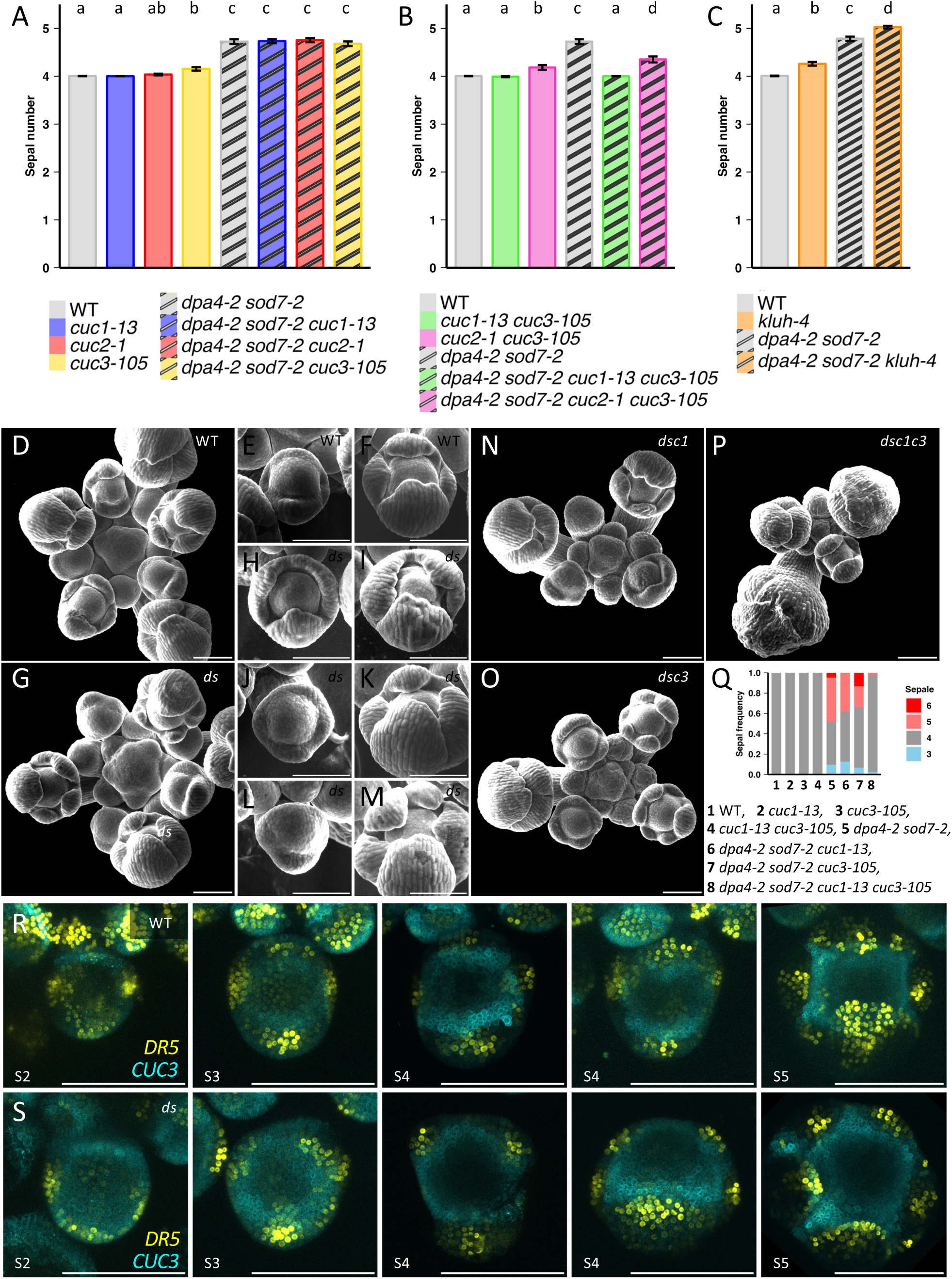
*DPA4*/*SOD7* control sepal patterning through the *CUC* genes and independently of *KLU.* **A.** Sepal number in WT, single *cuc*, *dpa4-2 sod7-2* double and *dpa4-2 sod7-2 cuc* triple mutants. **B.** Sepal number in WT, *cuc* double, *dpa4-2 sod7* double and *dpa4-2 sod7-2 cuc* quadruple mutants **C.** Sepal number in WT, single *klu-4*, *dpa4-2 sod7-2* double and *dpa4-2 sod7-2 klu-4* triple mutants. **D-P.** Scanning electron microscopy view of inflorescence apices or young developing floral meristems of wild-type (WT), *dpa4-2 sod7-2* (*ds*), *dpa4-2 sod7-2 cuc1-13* (*dsc1*), *dpa4-2 sod7-2 cuc3-105* (*dsc3*), *klu-4* (*k*) or *dpa4-2 sod7-2 2 cuc1-13 cuc3-105* (*dsc1c3*) plants. **Q.** Frequency of the flowers with different sepal primordia numbers in different genotypes **R,S.** Expression pattern in young developing flowers of pCUC3:CFP (CUC3 in cyan) and DR5:VENUS (DR5 in yellow) reporters in wild type (WT) and *dpa4-2 sod7-2* (*ds*). The floral stage is indicated (S2 to S5). The adaxial part of the floral meristem is always on the top of the image. In **A**-**C,** a Kruskal-Wallis test followed by a Dunn’s post-hoc test were performed to show significant differences between our different mutants (p- value<0.05). Scale bars = 100 µm in all panels.

The interaction between *DPA4*/*SOD7* and *CUC* genes for the control of petal number was more complex (Figure S3A,B,G,H), with for instance a reduction of the petal number below 4 in all three *dpa4-2 sod7-2 cuc* triple mutants. However, like for the sepals, the *dpa4-2 sod7-2 cuc1-13 cuc3-105* quadruple mutant showed an almost fully restored number of petals, with a mean petal number close to 4 and strongly reduced variability of the petal number (Figure S3B,H). Together, this suggested that the regulation of sepal and petal number by *DPA4* and*SOD7* occurs through a pathway involving *CUC1* and *CUC3*.

Like *ds,* the *klu-4* mutant showed an increase in sepal number, and both effects were additive as the sepal number further increased in the *dpa4-2 sod7-2 klu-4* triple mutant compared to *ds* and klu-4 (Figure 4C, Figure S3C,F,I). Petal number was also higher in *dpa4-2 sod7-2 klu-4* compared to *ds* and *klu-4.* This suggested that *DPA4*/*SOD7* and *KLU* act in independent pathways during floral organ specification (Figure 6D).

To further support the genetic interactions between the *NGAL*/*CUC*/*KLU* genes during flower development, we precisely characterized their expression using lines expressing simultaneously combinations of two transcriptional reporters of these genes (Figure S4). The five genes showed overall overlapping expression patterns in the boundary domain of inflorescence and floral meristems, with *KLU* being additionally expressed within the meristems. At floral stages S1 and S2 (floral stages defined according to Smyth et al. 1990), *DPA4* and *SOD7* reporters showed a strong expression in the adaxial domain, which was also observed for *CUC3* and to a lower extent *KLU* but not *CUC2*. In addition, *DPA4* and *CUC3* reporter expressions transiently extended into the entire floral meristem during S1. At S3, when the sepal-meristem boundary became marked by *CUC2* and *CUC3* reporter expressions, *DPA4* and *SOD7* reporter expression were not observed in the floral meristem. This domain was marked by *DPA4* and *SOD7* reporters at later stage S4, as were more inner boundary domains at S5. No expression was however observed for *DPA4* and *SOD7* in the intra whorl sepal-sepal boundary domain in contrast to *CUC2*. The outer abaxial epidermis of the sepal also strongly expressed the *SOD7* reporter. At S2 and S3, *KLU* was expressed throughout the floral meristem, with a stronger expression at its boundary with the inflorescence meristem at S2 and the sepal boundaries at S3. A strong expression of KLU remained in the center of the flower meristem at S4 and S5 and the sepal primordia maintained KLU expression at these stages. Together, these observations suggested that the genetic interaction between *DPA4*/*SOD7* and *CUC2*/*CUC3* genes occurs at the organ boundaries while the *KLU* effect on organ number independent of *DPA4*/*SOD7* may result from the unique expression of *KLU* at the meristem center.

### Interaction of *DPA4* and *SOD7* with *CUC* genes controls sepal initiation patterns

To further relate the changes in sepal numbers in mature flowers with early flower development we examined it by SEM (Figure 4D-P). While in the WT the stereotypical pattern of 4 initiating sepal primordia was observed (Figure 4D-F,Q), *ds* flowers showed highly variable patterns (Figure 4G-M,Q). Thus, we observed flowers with 3 equidistant and similar sepal primordia (Figure 4H,I) or flowers with 4 sepal primordia of inequal sizes and irregular positions (Figure 4J,K). Young flower primordia in which initiation of the adaxial sepal primordium appeared delayed could be observed (Figure 4L). Older flowers with 5 sepal primordia, including two small ones in adaxial position that could result from delayed initiation were present in the *ds* mutant (Figure 4M). Inactivating *CUC1* or *CUC3* alone in the *ds* background only very partially corrected sepal initiation defects, while inactivating *CUC1* and *CUC3* together almost totally suppressed *ds* sepal initiation defects (Figure 4N- Q).

To follow sepal initiation at the molecular level and relate it to *CUC* gene expression patterns, we imaged WT and *ds* young flowers expressing the auxin response marker *pDR5:VENUS* and *pCUC3:CFP* (Figure 4R,S). The *pDR5:VENUS* marker is expressed in the incipient organ primordia (Chandler et al. 2011; Kong et al. 2024). From floral stage S3 onwards, WT floral meristem had 4 separated DR5 expressing foci of cells, that correspond to the 4 sepal primordia initiation sites. By contrast, at S3 and S4, only 3 foci were often observed in *ds* flowers, one in the abaxial position, and two at an intermediate position between the median and adaxial positions. At later stages, five *pDR5:VENUS*- expressing cell foci were often observed, in agreement with the most frequent sepal primordium number observed in the *ds* mutants. At S2 stage, *pCUC3:CFP* expression was weak in WT floral meristems, while it appeared higher in *ds* floral meristems, in particular in its adaxial domain. This is also the region in which *pDPA4:GFP* and *pSOD7:GFP* are strongly expressed (Figure S4). Together, this suggest a model in which in the *ds* floral meristems, *CUC3* is transiently locally derepressed in the adaxial domain, which in turn interferes with auxin signaling and sepal primordium formation. As our genetic data indicate that *CUC1* is required with *CUC3* to cause abnormal sepal numbers in *ds,* we can speculate that *CUC1* may be similarly derepressed in *ds* and that it may also contribute to delayed sepal initiation as recently shown in the *drmy1* mutant (Kong et al. 2024).

### *DPA4/SOD7* and *CUC* genes independently repress seed growth

Mutation of the *KLU* gene leads to smaller seeds (Adamski et al. 2009) and totally suppresses the increase in seed size of *ds* double mutants (Zhang et al. 2015). Mutation of single *CUC* genes did not modify seed size in the *ds* mutant, but we observed a limited increase of seed size in single *cuc* mutants, which was however only statistically significant for *cuc3-105* (Figure 5A). This prompted us to analyze the effects of multiple *CUC* gene mutations on seed size to overcome possible genetic redundancy. Indeed, seeds of both *cuc1-13 cuc3-105* and *cuc2-1 cuc3-105* double mutant were larger than WT (Figure 5B, C): seeds of the third double mutant, *cuc1 cuc2,* could not be analyzed as this double mutant does not produce any flowers (Aida et al. 1997). A further increase in seed size was observed in the *ds* background when two *CUC* genes were simultaneously inactivated (Figure 5B,C). These observations indicate that *CUC* genes redundantly repress seed growth and do so independently of *DPA4* and *SOD7*. Again, as during cauline leaf development, our genetic analysis did not allow us to determine whether *CUC* genes act by repressing *KLU* or via a pathway independent of *KLU*. However, because *CUC* genes have been shown to activate *KLU* expression in embryos and leaves (Maugarny-Calès et al. 2019; Aida et al. 2020), we rather favor the hypothesis of an action of *CUC* genes on seed growth independent of *KLU* (Figure 6D). This model for the genetic interactions directing seed size regulation contrasts with the ones identified in other organs (Figure 6A-C), since here *DPA4*/*SOD7* appears to act independently but with similar effects as the *CUC* genes, while in other organs *DPA4*/*SOD7* and *CUC* genes act antagonistically. This prompted us to question whether *DPA4*/*SOD7* represses *CUC* genes during seed development as it does in other organs (Figure 3E, Engelhorn et al. 2012; Shao et al. 2020; Nicolas et al. 2022). To test this, we first questioned whether the expression of these genes overlap and, because *DPA4*/*SOD7* control seed size maternally by affecting ovule growth (Zhang et al. 2015), we analyzed their expression patterns during ovule development. *pDPA4:GFP is* strongly expressed in the outer cells of the basal part of the nucellus and weakly in the inner integument primordia in stage 2-III ovules (stages according to Schneitz et al., 1995) (Figure S5A). At later stages, *pDPA4:GFP* expression remained high at the base of the inner integuments but extended into the inner integuments and became expressed at the base of the outer integuments too (Figure S5B-D). The *pSOD7:GFP* reporter showed an expression pattern very close to the one provided by the *pDPA4:GFP* reporter (Figure S5E-H) (Zhang et al. 2015). *pKLU:GFP* reporter was expressed in domains that were also very similar to the ones of *pDPA4:GFP* and *pSOD7:GFP* (Figure S5M-P) (Adamski et al. 2009). In contrast, *pCUC2:RFP* and *pCUC3:CFP* showed very different patterns (Figure S5I-L). At stage 2-III, *pCUC3:CFP* was not detectable and *pCUC2:RFP* was limited to a few cells at the very base of the nucellus (Figure S5I). At stage 3-I and later, *pCUC3:CFP* formed a ring-link expression pattern at the junction of the nucellus and the inner integument, while *pCUC2:RFP* was expressed below (Figure S5J-L). Therefore, despite having different patterns, the *pCUC2:RFP* and *pCUC3:CFP* domains were included in the domains expressing the *pDPA4:GFP* and *pSOD7:GFP* reporters, raising the possibility that *DPA4*/*SOD7* could regulate *CUC* gene expression during ovule development. To test this, we analyzed *pCUC2:RFP* and *pCUC3:CFP* expression during *ds* and WT ovule development. At stages when integuments were initiated, expression of *pCUC2:RFP* was extended in the nucellus of *ds* ovules compared to WT (Figure 5D, E). At later stages, a higher and enlarged *pCUC2:RFP* expression was observed in *ds* ovules (Figure 5F, G). The expression of *pCUC3:CFP* was not modified in *ds* ovules (Figure 5D-G). Together, these data suggested that *DPA4*/*SOD7* repressed *CUC2* expression during ovule development, but that derepression of *CUC2* in *ds* was not causal for the increase in seed size of the double mutant.

**Figure 5.**
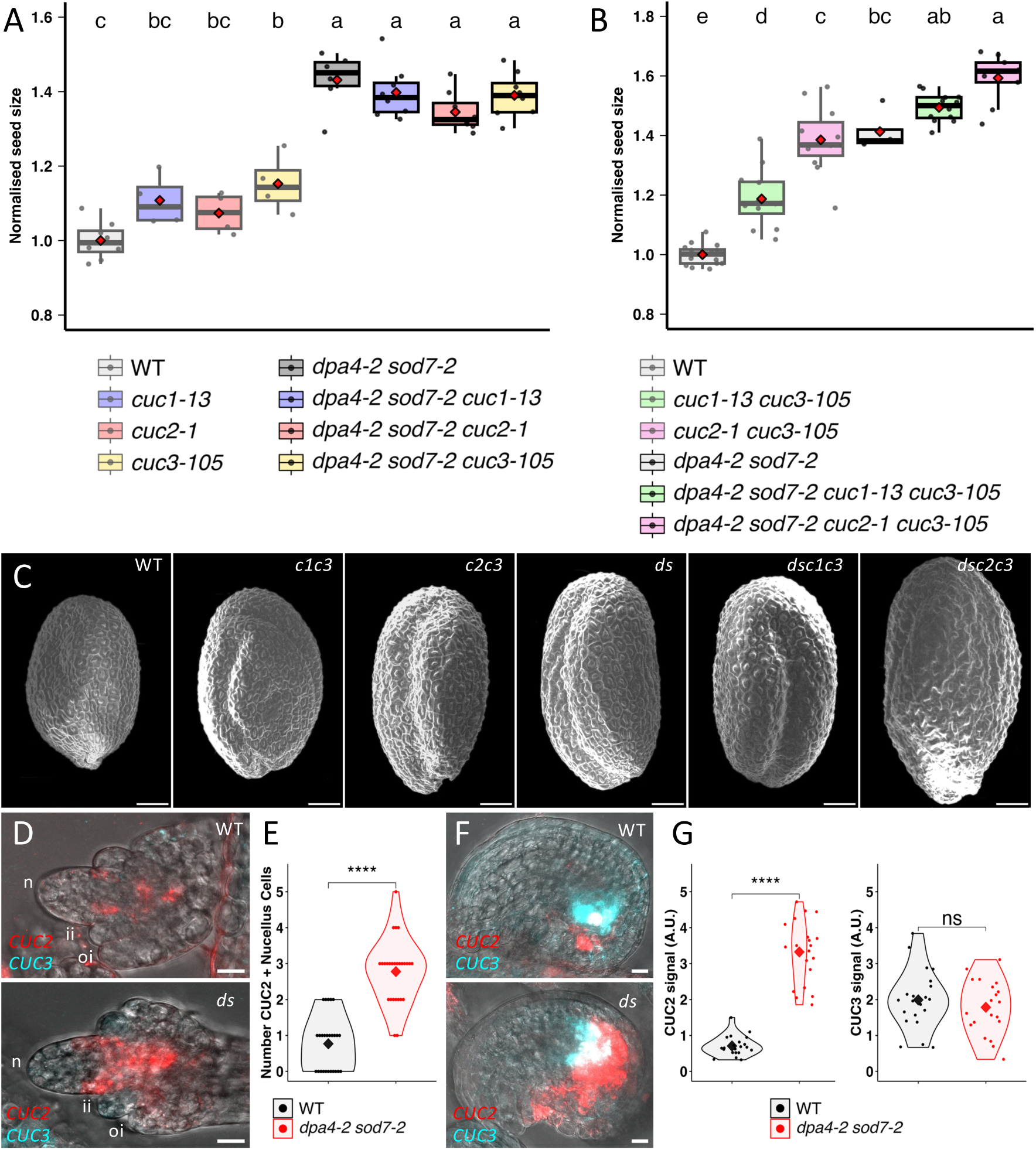
*DPA4*/*SOD7* and *CUC* genes additively repress seed growth. **A.** Seed size in WT, single *cuc*, *dpa4-2 sod7-2* double and *dpa4-2 sod7-2 cuc* triple mutants. **B.** Seed size in WT, *cuc* double, *dpa4-2 sod7* double and *dpa4-2 sod7-2 cuc* quadruple mutants **C.** Scanning electron microscopy view of wild-type (WT), *cuc1-13 cuc3-105* (*c1c3*), *cuc2-1 cuc3-105* (*c2c3*), *dpa4-2 sod7-2* (*ds*), *dpa4-2 sod7-2 cuc1-13 cuc3-105* (*dsc1c3*) and *dpa4-2 sod7-2 cuc2-1 cuc3-105* (*dsc2c3*) seeds **D.** Expression of *pCUC2:RFP* (in red) and *pCUC3:CFP* (in cyan) in developing ovules at stage 2-IV (Schneitz et al. 1995) of wild-type (WT) or *dpa4-2 sod7-2* double mutant (*ds*) plants.ii: inner integument, io: outer integument, n: nucellus **E.** Quantification of nucellus cells expressing *pCUC2:RFP.* Ovules were at stages2-III or 2-IV. **F.** Expression of *pCUC2:RFP* (in red) and *pCUC3:CFP* (in cyan) in developing ovules at stage 3-III (Schneitz et al. 1995) of wild-type (WT) or *dpa4-2 sod7-2* double mutant (*ds*) **G.** Quantification of *pCUC2:RFP* and *pCUC3:CFP* expression levels in ovules at stages 3-III to 3-V. In **A** and **B**, data were normalized to the mean value of the WT. Rectangle represent the first and third quartiles, with a horizontal central line marking the median. Mean values are represented as red diamonds. In **G**, statistical significance is tested by Student’s tests: p-value is 0.001 for ****. In **A** and **B,** ANOVA followed by a Tukey post-hoc test was performed (p-value<0.05). In **E,** statistical significance is tested by Mann & Whitney tests: p-value is 0.001 for **** Scale bars = 100 µm in C, 10 µm in D and F.

## Discussion

How gene regulatory networks (GRNs) centered around pleiotropic genes are remodeled remains a largely open question in developmental and evolutionary biology. In this study, we explored this issue by focusing on the NGAL family of transcription factors, which play pleiotropic roles in plant development. Hence, we determined the gene network linking *NGALs* with their downstream genes *KLU* and *CUCs* during the regulation of seven different biological processes (growth of cauline leaves, petals, pistils, seeds, initiation of sepals and petals, inflorescence phyllotaxis). We identified four distinct types of network organizations, each differing in how *NGAL* effects are mediated through either *CUCs*, *KLU*, or a combination of both (Figure 6D). The network regulating seed growth was notably distinct from the others. In this case, *NGAL* and *CUC* genes had additive effects on seed growth repression. In contrast, in the three other networks, *CUC* genes had either antagonistic roles to the *NGAL* genes or did not interact with the *NGAL* pathway. The network’s precise structure can be further refined due to genetic redundancy among the three *CUC* genes, with varying degrees of involvement for these different members. For instance, our data suggest that *CUC2* and *CUC3* had similar contribution to the *NGAL*-mediated regulation of cauline leaf growth while *CUC1* had no apparent role in it. Importantly, the structure of the network was not conserved within a specific organ or in the regulation of the specific biological process across different organs. For instance, we observed different genetic networks governing petal initiation and their subsequent growth. Similarly, the structures of the network governing the growth of cauline leaves, petals or seeds were also different, illustrating inter-organ variation.

**Figure 6.**
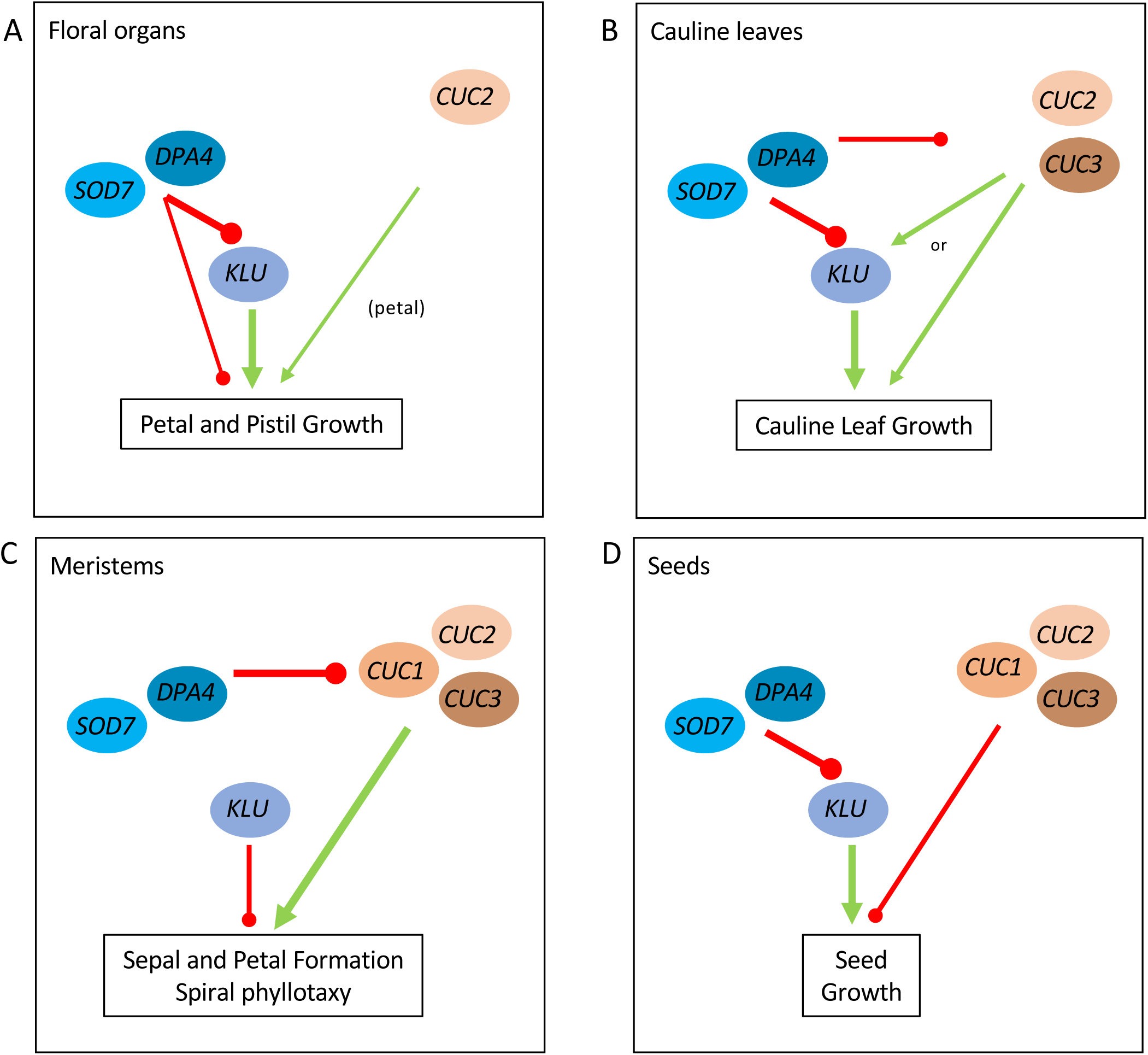
Organ-wide rewiring of the *NGAL*/*CUC*/*KLU* regulatory network. The GRNs were genetically determined by analyzing the phenotypes of single and multiple mutants. Gene expression patterns could be further used to support the GRNs. Repressing activities are shown by red lines ending by a point, while promoting activities are indicated by green arrows. Thickness of the red and green lines represent the importance of the regulatory step to the regulated biological process. **A.** The major pathway by which *DPA4*/*SOD7* repress petal and pistil growth acts through the repression of *KLU* pathway, which promotes growth of these organs. *DPA4*/*SOD7* also represses petal and pistil growth *via* a second, minor pathway independent of *KLU*. *DPA4*/*SOD7* act independently of the *CUC* genes. However, *CUC2* has a minor role in the promotion of petal growth. **B.** *DPA4*/*SOD7* repress cauline leaf growth via two pathways. The major pathway mediating the growth-repressing effect of *DPA4*/*SOD7* involves the repression of the *KLU* pathway, which promotes growth. The second, more minor, pathway operates through *CUC2* and *CUC3*, which may act either *via* or independently of the *KLU* pathway. **C.** *DPA4*/*SOD7* determine the patterns of phyllotaxy and sepal and petal initiation *via* a pathway involving the *CUC* genes. Contribution of the individual *CUC* genes to this varies (not shown in the figure). For instance, *CUC3* has an important contribution to phyllotaxy while *CUC1* has a more minor role. In contrast, both *CUC1* and *CUC3* have an important contribution to sepal patterning. The *KLU* pathway has also a minor contribution in the determination of the sepal and petal initiation patterns and phyllotaxy, this role is however independent of *DPA4*/*SOD7* pathway. **D.** *DPA4*/*SOD7* repress seed growth *via* the *KLU* pathway (Zhang et al. 2015). The *CUC* genes also repress seed growth. *DPA4*/*SOD7*-mediated seed growth repression is independent of the *CUC* genes, although *DPA4*/*SOD7* represses *CUC2* expression during ovule development (not shown in the figure).

To establish these gene networks, we used morphometric analyses across various genetic backgrounds, combined with gene expression characterization using imaging of reporter constructs. These methods are particularly well-suited for analyzing inter organ variations, as they allow linking functional roles with spatial resolution of gene expression domains. Alternative methods can be used to establish GRNs at the molecular level with different spatial resolutions (O’Malley et al. 2016; Qian and Huang 2020; Tripathi and Wilkins 2021; Li et al. 2023; Yin et al. 2023; Yuan and Duren 2024). Combining spatial information provided by single cell analysis or other cell-type specific approaches helps resolving GRNs at the organ or cell type level (Baumgart et al.; Brady et al. 2011; Tian et al. 2014; de Luis Balaguer et al. 2017; Sonawane et al. 2017; Kajala et al. 2021; Robertson and Wilkins 2023). Such approaches have also been used to follow evolution of GRNs during cell differentiation or in response to pathogens (Shahan et al. 2022; Kamimoto et al. 2023; Nobori et al. 2023; Zhang et al. 2023; Zhu et al. 2023) or to compare GRN evolution across organs or species (Kajala et al. 2021; Guillotin et al. 2023). However, they do not necessarily establish the functional roles of different network elements. Indeed, our study provides an example where a regulatory node conserved at the molecular level is functionally relevant in some organs but not in all. For instance, *NGAL* repression of *CUC2* contributes to the regulation of phyllotaxis, floral organ initiation, and cauline leaf growth, but not seed growth. More specifically, *DPA4*/*SOD7* repression of *CUC2* transcription occurs during seed development but does not seem to contribute to larger *ds* seeds, as the *dpa4 sod7* mutant does not show any restoration of seed size when *CUC2* is mutated. These observations suggest that GRNs can be explored at both molecular and functional levels, and that these two levels may not always perfectly align.

Several hypotheses may explain why a regulation occurring at the molecular level has no apparent functional effect as we found for *CUC2* transcriptional repression by *NGAL* genes during ovule development. First, gene regulation occurs at multiple levels, and in the case of *CUC2*, post-transcriptional regulation by the microRNA miR164 and protein-protein interactions are important factors regulating its activity (Nikovics et al. 2006; Rubio-Somoza et al. 2014; Barro-Trastoy et al. 2022) and miR164 can even mask the effects of a transcriptional derepression (Maugarny et al. 2024). A second, more specific hypothesis regarding *CUC* genes arises from their complex effects on growth. Indeed, *CUC2* represses growth autonomously, in part by modifying the expression of cell wall modifying enzymes (Peaucelle et al. 2007; Kawamura et al. 2010; Kierzkowski et al. 2019; Serra and Perrot-Rechenmann 2020; Bouré et al. 2022), while it promotes growth at a distance in part via the modulation of auxin response (Shankar et al.; Bilsborough et al. 2011; Maugarny-Calès et al. 2019; Hu et al. 2024). Hence, predicting the effects on growth of *CUC* modulation may be difficult, and for instance both larger or smaller leaves have been shown to result from *CUC2* over expression, possibly as a result from different pattern and/or level of expression (Nikovics et al. 2006; Larue et al. 2009; Li et al. 2020). Finally, and more specifically concerning seed size, both maternal and zygotic factors play a role in its regulation (Li et al. 2024). Here we concentrated on the maternal control, as *NGAL* genes were shown to affect seed size through this pathway (Zhang et al. 2015). However, the *NGAL*/*KLU*/*CUC* module which is also expressed during embryo development (Aida et al. 1999, 2020; Nicolas and Laufs 2022) may contribute to the regulation of seed size via the zygotic pathway, a hypothesis that remains to be tested.

Our work identifies the modulation of gene expression patterns as an important mechanism leading to inter-organ changes in the *NGAL*/*CUC*/*KLU* GRN. This is supported by the comparison of growth regulation between petals and cauline leaves. Indeed, genetic analysis indicate that *NGAL* repression of growth is in part mediated by the *CUC2* and *CUC3* genes in cauline leaves, while it is independent from these genes in petals. We could relate this difference to the expression of the *CUC2* and *CUC3* genes at the cauline leaf margin where their expression overlaps with that of *DPA4* and *SOD7*, while the expression of the *CUC* genes is absent from the petal margin. Although leaves and petals are related organs, they also differ in their developmental and growth programs (Pelaz et al. 2001; Sauret-Güeto et al. 2013; Le Gloanec et al. 2024). Indeed, we observed different patterns of auxin response distribution between petals and cauline leaves, which could contribute to the difference in *CUC* gene expression between these organs (Hay et al. 2006; Kawamura et al. 2010; Bilsborough et al. 2011). Together, this suggests that developmental constraints, by influencing gene expression patterns, may contribute to inter-organ modulation of GRNs. However, variations in gene expression patterns can also influence GRNs through an alternative mechanism, as demonstrated in sepal initiation. In this second case, genetic and expression data converge to support a model in which sepal and petal primordia numbers are regulated by two different independent pathways: the first acting at the organ boundary involves *DPA4*/*SOD7* repressing *CUC2*/*CUC3* and the second acting more at the meristem center involves *KLU*. In this model, *KLU* activity at the meristem center would escape *DPA4*/*SOD7* regulation by being expressed in a domain where *DPA4*/*SOD7* expression is absent. These two examples illustrate that variations in the combinatorial expression of the GRN actors are a major driver of GRN rewiring.

Here, we show that the pleiotropic effects of *NGAL* genes are linked to an organ-wide reconfiguration of their associated GRN, involving the *KLU* and *CUC* genes. The *KLU* and *CUC* pathways have distinct biological functions: *KLU* is primarily involved as a growth regulator (Anastasiou et al. 2007; Adamski et al. 2009; Eriksson et al. 2010; Kazama et al. 2010; Chakrabarti et al. 2013; Zhang et al. 2015), whereas *CUC* genes not only influence growth but also play a crucial role in establishing developmental patterns (Furutani et al. 2004; Bilsborough et al. 2011; Maugarny et al. 2016; Maugarny-Calès et al. 2019). Therefore, the rewiring of the GRN provides as a mechanism to balance the relative contributions of the *KLU* and *CUC* pathways across different organs, thereby determining the GRN’s impact on growth or patterning. In this view, GRN rewiring represents a means to shape diverse biological outcomes. More generally, we propose that organ-wide GRN plasticity facilitates the pleiotropic roles of genes.

## Materials and Methods

### Plant materials

The *ngal* and *cuc* single and multiple mutants (Nicolas et al. 2022) and *klu-4* mutant (Anastasiou et al. 2007) and the *pKLU:GFP, pDR5:VENUS*, (Maugarny-Calès et al. 2019), *pCUC2:RFP*, *pCUC3:CFP*, *pDPA4:GFP* and *pSOD7:GFP* (Nicolas et al. 2022) reporter lines have been described before. New combinations of mutants or mutants expressing reporter lines were generated by crossing. Lines homozygous for the desired mutation(s) were identified by PCR-based genotyping. The presence of the reporter line was verified through its expression and its homozygous state was determined by analyzing its segregation in the next generation. Only lines homozygous for the reporter(s) were analyzed except for the comparison between WT and *dpa4-2 sod7-2* of *pCUC2:RFP* and *pCUC3:CFP* (Figures 3E, 5D,F) and the combinations of two reporters (Figure 4R,S, Figure S4A-E, G,H), in which each reporter was at the heterozygous stage.

### Plant Growth and Phenotyping

Plants were grown in the greenhouse in a mix peat/sand (TREF 1018201170, TREF, France). The length of the 3 first (most basal) cauline leaves was measured using a ruler on plants grown in the greenhouse. Floral organ numbers were determined under a binocular. Petal and pistils were dissected from mature flowers, mounted in water, imaged using a binocular and their size was measured using FIJI (Schindelin et al. 2012). For internode length, fully grown inflorescences were taped on paper, scanned and the distances between successive nodes were measured using ImageJ. Flowers were defined as belonging to clusters when they were separated by internodes shorter than 3 mm. For seed size, seeds were sprayed on tape, imaged using a scanner and seed area was measured using a dedicated macro on ImageJ (Seed XXXXX). Images were threshold and the area of individual particles was measured. To remove possible non-seed particle, first size (> 0.05 mm^2^) and circularity (0.75-1.00) filters were applied, the result of this filtering was then inspected manually. In particular, we removed data from particles which were fused, such fused seeds as described previously for *cuc1 cuc3* and *cuc2 cuc3* double mutants (Gonçalves et al. 2015). Each point in the graph represents the mean area of at least 75 seeds of one plant.

### Data representation and statistical analysis

All data were analysed using R (R Core Team 2022). Boxplots (for cauline leaf, petal lengths, pistil width and seed size), were generated using the geom_boxplot function of the ggplot2 package (Wickham 2016). They compactly display the distribution of a continuous variable by allowing the visualization of the median (bold horizontal bar), the first and third quartile (respectively lower and upper hinges), the lowest and largest values no further than 1.5 * interquartile range from the hinge (respectively lower and upper whisker). In addition, all data are represented as individual points and the mean is represented. Values were normalized relative to the mean value of the WT. In bar plots (for floral organ numbers) error bars represent the standard errors. Violin plots (for internode length) were generated using the geom_violin function (ggplot2 package). Horizontal black bars represent the median of the values.

Statistically significant differences were shown using an ANOVA followed by Tukey’s post-hoc test (p-value <0.05) (Figure 1C, I; 5A, B), a nonparametric Kruskal-Wallis test followed by Dunn’s post-hoc test (p-value <0.05) (Figure 1A, E, G; S1A-D, I; 2B-E; S2B, C; S3A-C; 4A-C), a Student’s t test (p-value < 0.05) (Figure 5G), or using a non-parametric Mann & Whitney test (p-value < 0.05) (Figure 5E). The application conditions of ANOVA or Student’s t test were checked using the Shapiro-Wilk (distribution) and Levene (homogeneity of variances) tests respectively. Comparison of the internode length distributions (Figure 4A, B) were done by a two sample Kolmogorov-Smirnov test (p-value <0.01). The R packages car, dplyr, FSA and rcompanion were used for statistical analysis. Letters were used to highlight the significant phenotypic differences between different genotypes.

### Confocal imaging and quantification

Confocal imaging was performed on a Leica SP5 inverted microscope (Leica Microsystems, Wetzlar, Germany). The lenses were Leica 20x or 40x HCX PL APO CS. Acquisition parameters are presented in Supplemental Table S1. Maximum intensity projections along the Z axis were made using ImageJ. Stitching of several cauline leaf pictures was performed using the Pairwise Stiching plugin (Preibisch et al. 2009) and FigureJ (Mutterer and Zinck 2013) was used to assemble the figures.

For the comparison of *pCUC2:RFP* and *pCUC3:CFP* expression in ovules imaging was performed using the confocal same settings (laser power and signal gain) for WT and *dpa4-*2 *sod7-2*. For signal quantification, we first identified the median Z-section (with the two spots of *pCUC3:CFP* expression most distant), next we performed a max intensity projection along the Z axis including the 5 sections below and above the median section (Z step =2 µm). Fluorescent signal was then measured in the entire ovule (excluding the funiculus) and the background was subtracted from it.

### Scanning electron microscopy

Freshly sampled tissues were cooled to −33C° by a Peltier cooling stage (Deben) and observed with a Hirox SH-1500 benchtop scanning electron microscope.

## Acknowledgments

The IJPB benefits from the support of Saclay Plant Sciences-SPS (ANR-17-EUR-0007). This work has benefited from the support of IJPB’s Plant Observatory platforms PO-Plants and PO-Cyto. PP was partly supported by a scholarship from the Erasmus+ program of the European Union. We thank Michael Lenhard for helpful comments on the manuscript.

## Author contributions

Conceptualization: AN, AM, PL

Formal Analysis: AN, PP, MC

Investigation: AN, PP, MC, BA, MG, NA, PL

Writing – Original Draft: PL

Writing – Review & Editing: AN, PP, MC, NA, AM, PL

Visualization: AN, PL

Supervision: PL, AM

---

**Supplemental Figure 1. Floral organ numbers and petal length in single and double *ngal* mutants**

**A-D.** Floral organ numbers in WT, single and double *ngal* mutants (**A**, sepal; **B**, petal; **C**, stamen; **D**, carpel). in single and double *ngal* mutants

**E-H.** Floral organ number frequency WT, in single and double *ngal* mutants (**E**, sepal; **F**, petal; **G**, stamen; **H**, carpel). The color coding of the organ number is specific for each organ type and shown below the graphs.

**I.** Petal length in WT, single and double *ngal* mutants. Data were normalized to the mean value of the WT.

In **A-D,** a Kruskal-Wallis test followed by a Dunn’s post-hoc test were performed to show significant differences between different mutants (p-value<0.05).

**Supplemental Figure 2. *DPA4*/*SOD7* redundantly repress pistil growth via *KLU* and auxin response in petal and cauline leaf.**

**A.** Scanning electron microscopy detail of wild type (WT), *dpa4-2 sod7-2* (*ds*), *cuc3-105* (*c3*), *dpa4-2 sod7-2 cuc3-105* (*dsc3*), *klu-4* (*k*) and *dpa4-2 sod7-2 klu-4* (*dsk*) pistil top part.

**B.** Ovary width in WT, single *cuc*, *dpa4-2 sod7-2* double and *dpa4-2 sod7-2 cuc* triple mutants.

**C.** Ovary width in WT, single *klu-4*, *dpa4-2 sod7-2* double and *dpa4-2 sod7-2 klu-4* triple mutants.

**D.** *DR5:VENUS* (*DR5*) expression pattern in yellow marking the auxin response in a young petal and cauline leaf. Chlorophyll fluorescence is shown in red. Scale bars = 100 µm in A and D.

In **B** and **C**, data were normalized to the mean value of the WT. Rectangle represent the first and third quartiles, with a horizontal central line marking the median. Mean values are represented as red diamonds.

In **B** and **C,** a Kruskal-Wallis test followed by a Dunn’s post-hoc test were performed to show significant differences between different mutants (p-value<0.05).

**Supplemental Figure 3. *DPA4*/*SOD7* control sepal and petal patterning via the *CUC* genes.**

**A.** Petal number in WT, single *cuc*, *dpa4-2 sod7-2* double and *dpa4-2 sod7-2 cuc* triple mutants.

**B.** Petal number in WT, *cuc* double, *dpa4-2 sod7* double and *dpa4-2 sod7-2 cuc* quadruple mutants

**C.** Petal number in WT, single *klu-4*, *dpa4-2 sod7-2* double and *dpa4-2 sod7-2 klu-4* triple mutants.

**D.** Sepal frequency in WT, single *cuc*, *dpa4-2 sod7-2* double and *dpa4-2 sod7-2 cuc* triple mutants.

**E.** Sepal frequency in WT, *cuc* double, *dpa4-2 sod7* double and *dpa4-2 sod7-2 cuc* quadruple mutants

**F.** Sepal frequency in WT, single *klu-4*, *dpa4-2 sod7-2* double and *dpa4-2 sod7-2 klu-4* triple mutants.

**G.** Petal frequency in WT, single *cuc*, *dpa4-2 sod7-2* double and *dpa4-2 sod7-2 cuc* triple mutants.

**H.** Petal frequency in WT, *cuc* double, *dpa4-2 sod7* double and *dpa4-2 sod7-2 cuc* quadruple mutants

**I.** Petal frequency in WT, single *klu-4*, *dpa4-2 sod7-2* double and *dpa4-2 sod7-2 klu-4* triple mutants.

In **A**-**C,** a Kruskal-Wallis test followed by a Dunn’s post-hoc test was performed to show significant differences between our different mutants (p-value<0.05).

**Supplemental Figure 4. *DPA4, SOD7, CUC2, CUC2* and *KLU* reporter expression during flower development.**

**A.** Inflorescence apex of a plant expressing simultaneously *pSOD7:GFP* (SOD7, green) and *pCUC2:RFP* (CUC2, red) reporters. The two first panels show separated channels while the last one shows the overlay.

**B.** Inflorescence apex of a plant expressing simultaneously *pDPA4:GFP* (DPA4, green) and *pCUC2:RFP* (CUC2, red) reporters. The two first panels show separated channels while the last one shows the overlay.

**C.** Inflorescence apex of a plant expressing simultaneously *pKLU:GFP* (KLU, green) and *pCUC3:CFP* (CUC3, cyan) reporters. The two first panels show separated channels while the last one shows the overlay.

**D to H**. Details of reporter expression at different floral stages indicated on the right (S1 to S5). The column **D** shows *pSOD7:GFP* (SOD7, green) expression. The column **E** shows *pCUC2:RFP* (CUC2, red) expression. The column **F** shows *pDPA4:GFP* (DPA4, green) expression. The column **G** shows *pKLU:GFP* (KLU, green) expression. The column **H** shows *pCUC3:CFP* (CUC3, cyan) expression. Note that *pSOD7:GFP* and *pCUC2:RFP* are expressed simultaneously in the same plants, as are *pKLU:GFP* and *pCUC3:CFP*.

Scale bars = 50 µm in all panels.

**Supplemental Figure 5. *DPA4, SOD7, CUC2, CUC2* and *KLU* reporter expression during ovule development.**

**A-D.** Developing ovules expressing a *pDPA4:GFP* (*DPA4*, green) reporter.

**E-H.** Developing ovules expressing a *pSOD7:GFP* (*SOD7*, green) reporter.

**E-H.** Developing ovules expressing simultaneously a *pCUC2:RFP* (*CUC2,* red) and *pCUC3:CFP* (*CUC3,* cyan) reporter.

**E-H.** Developing ovules expressing a *pKLU:GFP* (*KLU*, green) reporter. Bars = 50 µm in all panels

## Notes

### Competing Interest Statement

The authors have declared no competing interest.

