## Supplemental Figures and Table for "Developmental rewiring of the *NGAL*/*CUC*/*KLU* network associated with pleiotropic roles of *NGAL* genes"

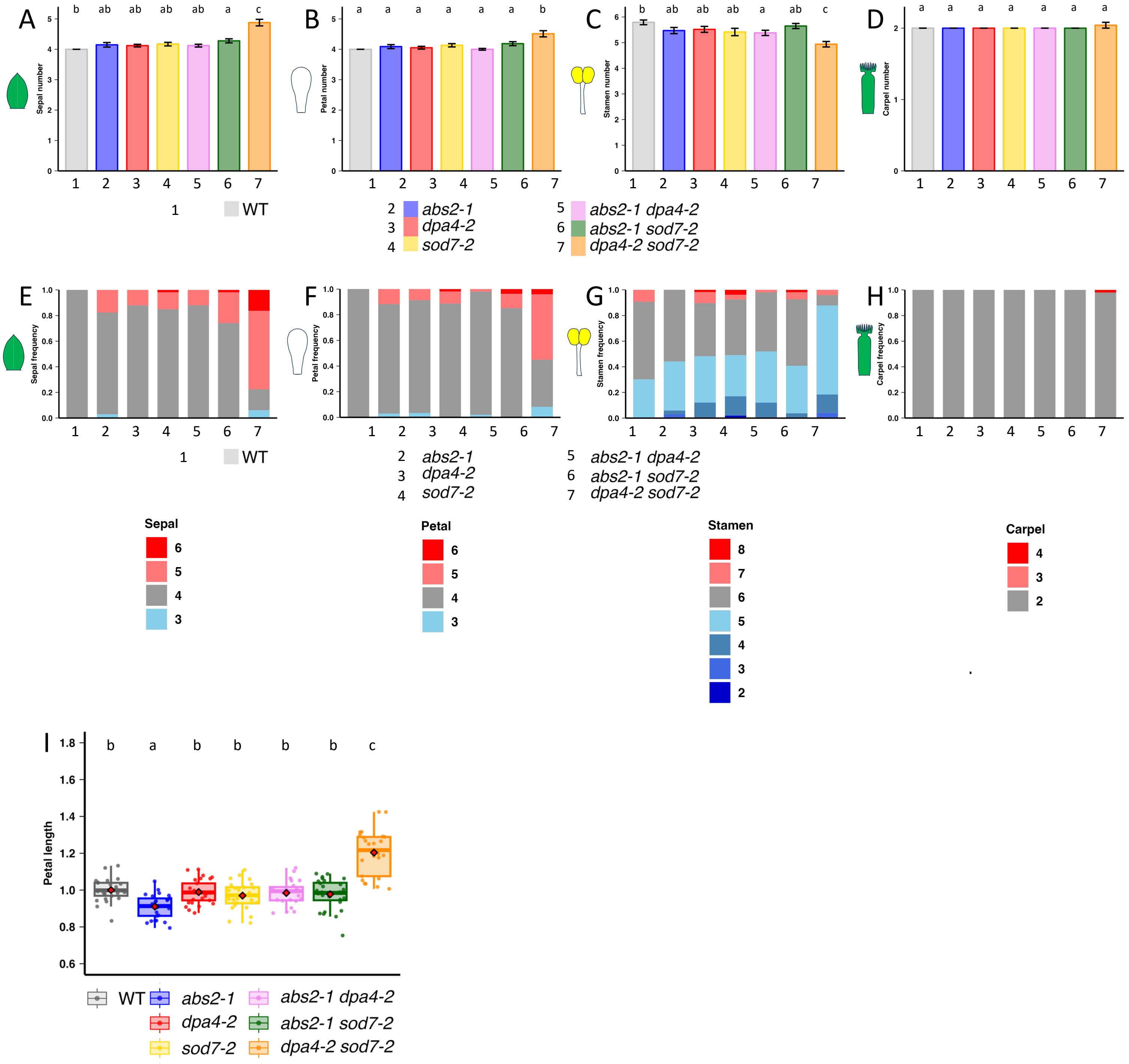


**Supplemental Figure 1. Floral organ numbers and petal length in single and double *ngal* mutants**

**A-D.** Floral organ numbers in WT, single and double *ngal* mutants (**A**, sepal; **B**, petal; **C**, stamen; **D**, carpel).

in single and double *ngal* mutants

**E-H.** Floral organ number frequency WT, in single and double *ngal* mutants (**E**, sepal; **F**, petal; **G**, stamen; **H**, carpel). The color coding of the organ number is specific for each organ type and shown below the graphs.

**I.** Petal length in WT, single and double *ngal* mutants. Data were normalized to the mean value of the WT.

In **A-D,** a Kruskal-Wallis test followed by a Dunn's post-hoc test were performed to show significant differences between different mutants (p-value<0.05).

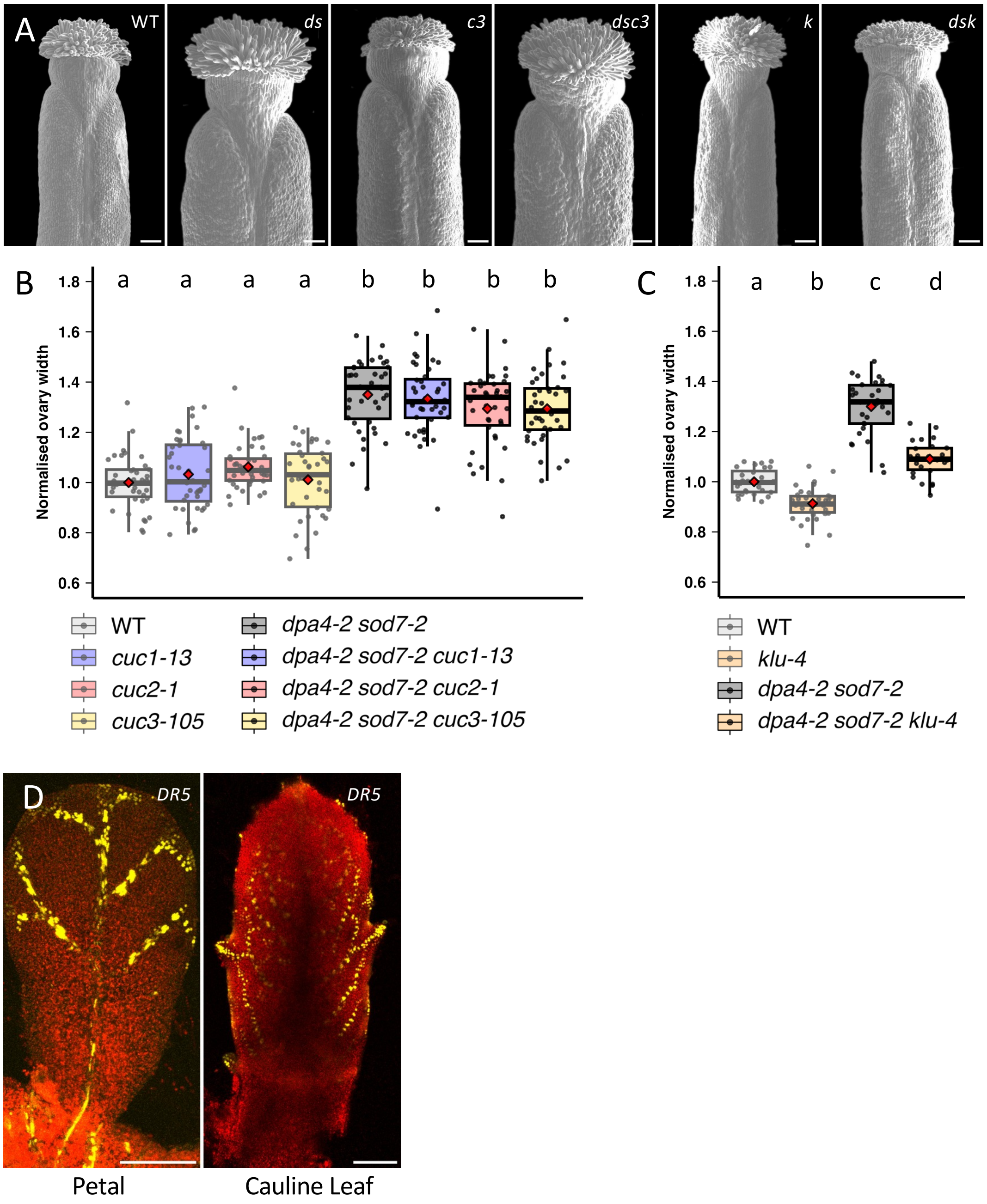


**Supplemental Figure 2. *DPA4/SOD7* redundantly repress pistil growth via *KLU* and auxin response in petal and cauline leaf.**

**A.** Scanning electron microscopy detail of wild type (WT), *dpa4-2 sod7-2* (*ds*), *cuc3-105* (*c3*), *dpa4-2 sod7-2 cuc3-105* (*dsc3*), *klu-4* (*k*) and *dpa4-2 sod7-2 klu-4* (*dsk*) pistil top part.

**B.** Ovary width in WT, single *cuc*, *dpa4-2 sod7-2* double and *dpa4-2 sod7-2 cuc* triple mutants.

**C.** Ovary width in WT, single *klu-4*, *dpa4-2 sod7-2* double and *dpa4-2 sod7-2 klu-4* triple mutants.

**D.** *DR5:VENUS* (*DR5*) expression pattern in yellow marking the auxin response in a young petal and cauline leaf. Chlorophyll fluorescence is shown in red. Scale bars = 100 µm in A and D.

In **B** and **C**, data were normalized to the mean value of the WT. Rectangle represent the first and third quartiles, with a horizontal central line marking the median. Mean values are represented as red diamonds.

In **B** and **C,** a Kruskal-Wallis test followed by a Dunn's post-hoc test were performed to show significant differences between different mutants (p-value<0.05).

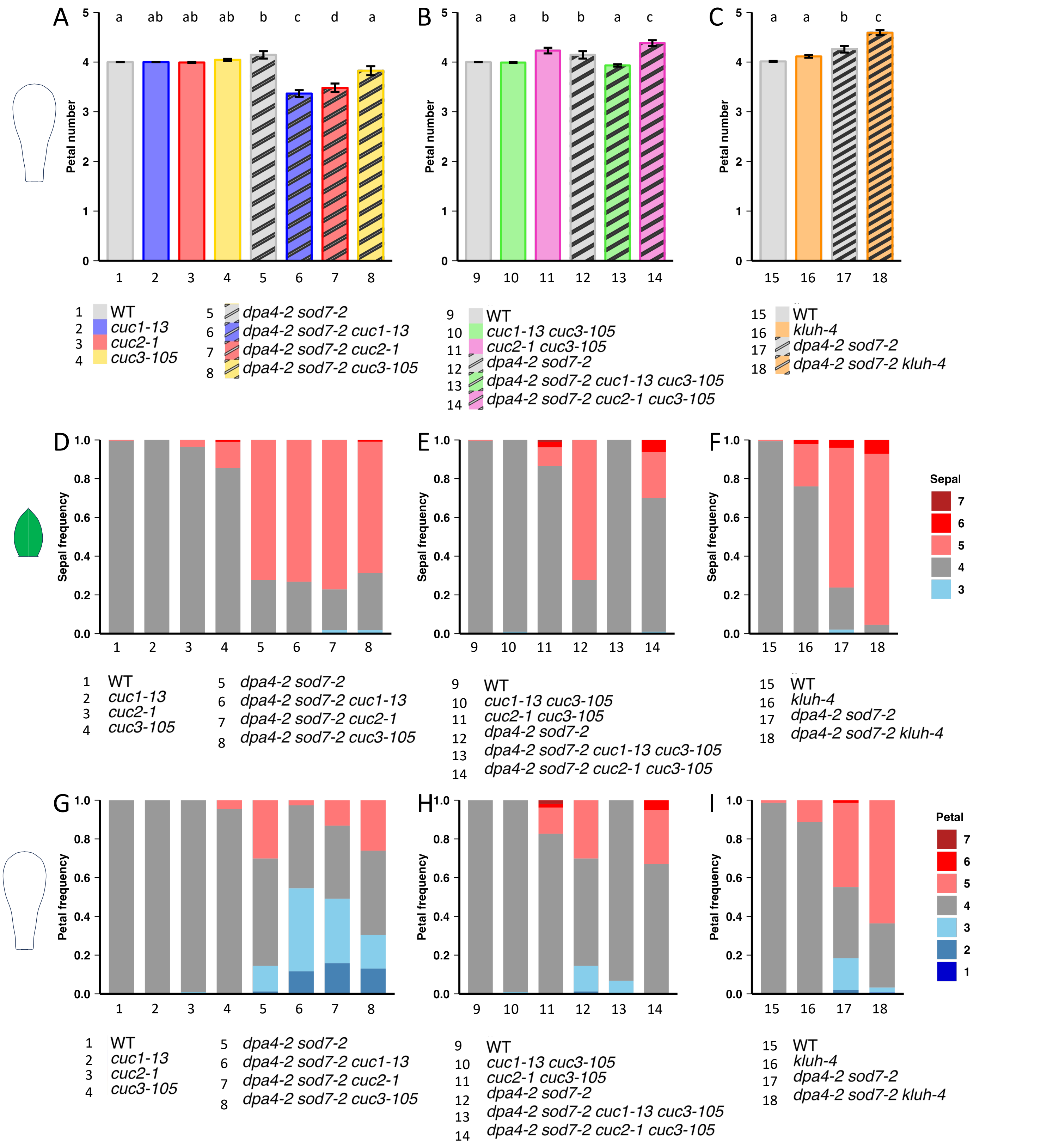


**Supplemental Figure 3. *DPA4/SOD7* control sepal and petal patterning via the *CUC* genes.**

**A.** Petal number in WT, single *cuc*, *dpa4-2 sod7-2* double and *dpa4-2 sod7-2 cuc* triple mutants.

**B.** Petal number in WT, *cuc* double, *dpa4-2 sod7* double and *dpa4-2 sod7-2 cuc* quadruple mutants

**C.** Petal number in WT, single *klu-4*, *dpa4-2 sod7-2* double and *dpa4-2 sod7-2 klu-4* triple mutants.

**D.** Sepal frequency in WT, single *cuc*, *dpa4-2 sod7-2* double and *dpa4-2 sod7-2 cuc* triple mutants.

**E.** Sepal frequency in WT, *cuc* double, *dpa4-2 sod7* double and *dpa4-2 sod7-2 cuc* quadruple mutants

**F.** Sepal frequency in WT, single *klu-4*, *dpa4-2 sod7-2* double and *dpa4-2 sod7-2 klu-4* triple mutants.

**G.** Petal frequency in WT, single *cuc*, *dpa4-2 sod7-2* double and *dpa4-2 sod7-2 cuc* triple mutants.

**H.** Petal frequency in WT, *cuc* double, *dpa4-2 sod7* double and *dpa4-2 sod7-2 cuc* quadruple mutants

**I.** Petal frequency in WT, single *klu-4*, *dpa4-2 sod7-2* double and *dpa4-2 sod7-2 klu-4* triple mutants.

In **A**-**C,** a Kruskal-Wallis test followed by a Dunn's post-hoc test was performed to show significant differences between our different mutants (p-value<0.05).

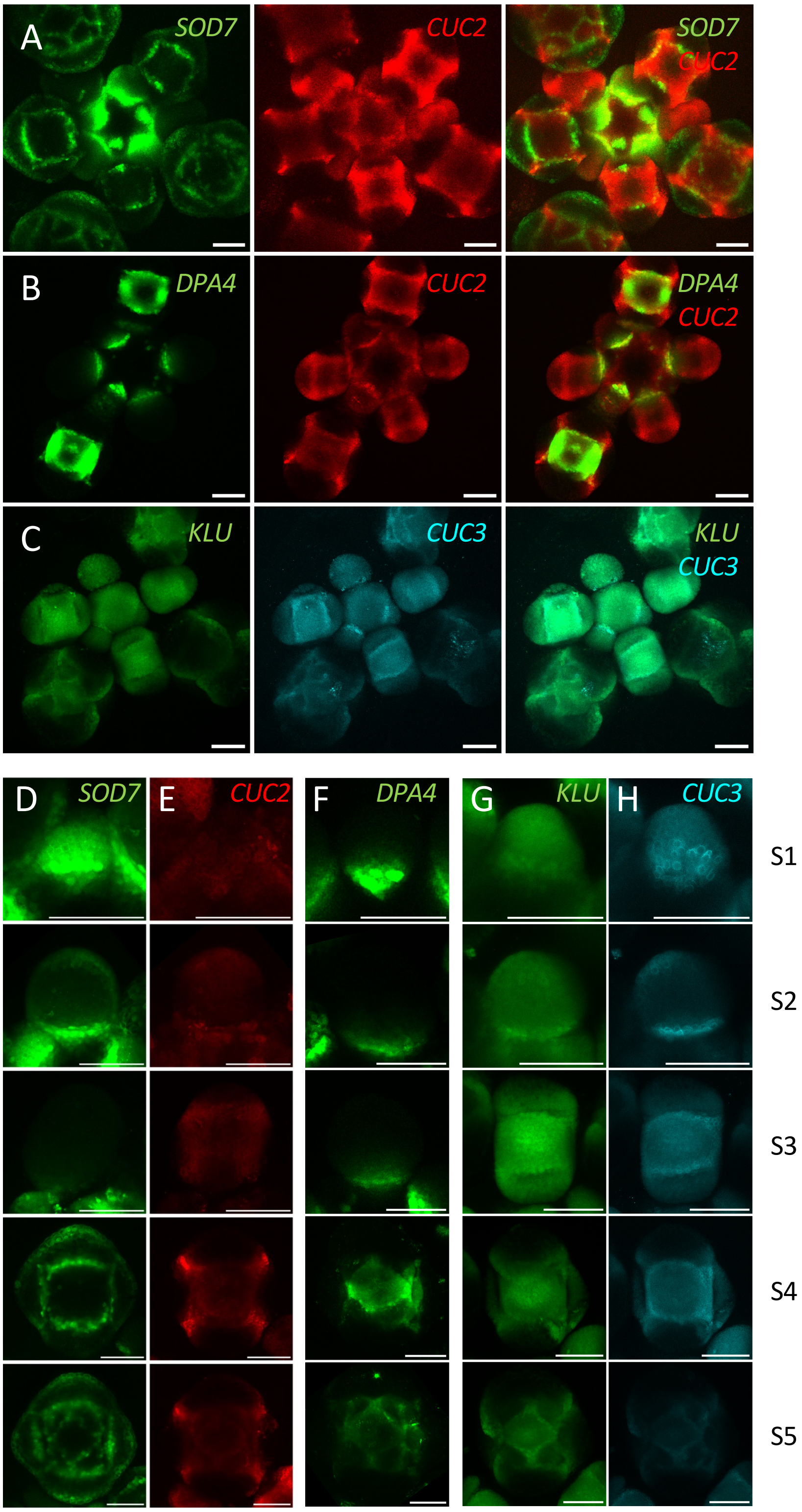


**Supplemental Figure 4. *DPA4, SOD7, CUC2, CUC2* and *KLU* reporter expression during flower development.**

**A.** Inflorescence apex of a plant expressing simultaneously *pSOD7:GFP* (SOD7, green) and *pCUC2:RFP* (CUC2, red) reporters. The two first panels show separated channels while the last one shows the overlay.

**B.** Inflorescence apex of a plant expressing simultaneously *pDPA4:GFP* (DPA4, green) and *pCUC2:RFP* (CUC2, red) reporters. The two first panels show separated channels while the last one shows the overlay.

**C.** Inflorescence apex of a plant expressing simultaneously *pKLU:GFP* (KLU, green) and *pCUC3:CFP* (CUC3, cyan) reporters. The two first panels show separated channels while the last one shows the overlay.

**D to H**. Details of reporter expression at different floral stages indicated on the right (S1 to S5).

The column **D** shows *pSOD7:GFP* (SOD7, green) expression. The column **E** shows *pCUC2:RFP* (CUC2, red) expression. The column **F** shows *pDPA4:GFP* (DPA4, green) expression. The column **G** shows *pKLU:GFP* (KLU, green) expression. The column **H** shows *pCUC3:CFP* (CUC3, cyan) expression. Note that *pSOD7:GFP* and *pCUC2:RFP* are expressed simultaneously in the same plants, as are *pKLU:GFP* and *pCUC3:CFP*.

Scale bars = 50 µm in all panels.

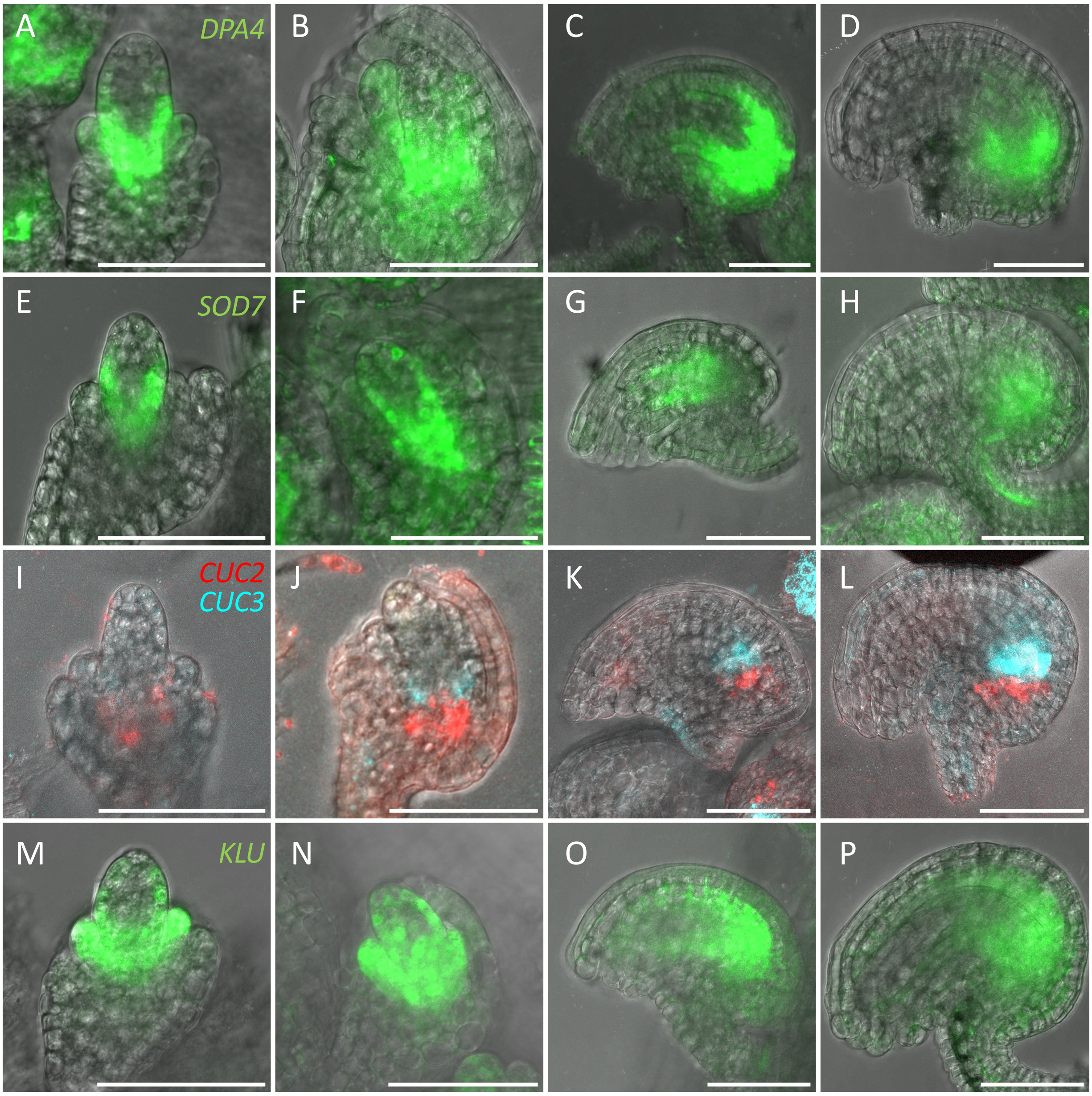


**Supplemental Figure 5. *DPA4, SOD7, CUC2, CUC2* and *KLU* reporter expression during ovule development.**

**A-D.** Developing ovules expressing a *pDPA4:GFP* (*DPA4*, green) reporter.

**E-H.** Developing ovules expressing a *pSOD7:GFP* (*SOD7*, green) reporter.

**E-H.** Developing ovules expressing simultaneously a *pCUC2:RFP* (*CUC2,* red) and *pCUC3:CFP* (*CUC3,* cyan) reporter.

**E-H.** Developing ovules expressing a *pKLU:GFP* (*KLU*, green) reporter.

Bars = 50 µm in all panels

| Reporter | Excitation wavelength (nm) | Detection wavelength (nm) |
|---|---|---|
| **pCUC2:RFP** | 561 | 590-631 |
| **pCUC3:CFP** | 458 | 470-512 |
| **pKLUH:GFP, pDPA4:GFP, pSOD7:GFP** | 488 | 498-535 |
| **pDR5:VENUS** | 514 | 520-555 |

**Table S1: Acquisition parameters for confocal reporter imaging**
